# Base Composition Influences the Position and Precision of RNA Polymerase II Disassociation in Basal and Perturbed Conditions

**DOI:** 10.64898/2025.12.02.691879

**Authors:** Georgia E.F. Barone, Jacob T. Stanley, Daniel Ramirez, Joseph F. Cardiello, Nina Ripin, Roy Parker, Mary A. Allen, Robin D. Dowell

## Abstract

RNA Polymerase II (Pol II) transcribes all protein-coding and many non-protein coding genes in the genome. Pol II transcription termination is crucial for mRNA maturation and, when disrupted, can lead to altered mRNA processing and mRNA export. Termination involves two intertwined processes: pre-mRNA cleavage and Pol II release from the DNA (disassociation). Despite its importance, the exact mechanisms underlying Pol II disassociation from the DNA remain poorly understood. Moreover, under certain cellular stress conditions, there is a partial failure of cleavage, leading to a shift of the position of disassociation further downstream, a phenomenon known as run-on transcription. We performed the first-ever systematic analysis of Pol II termination across cell types and species and provide novel insights into the mechanism of disassociation. Using a probabilistic mixture model to quantify Poll II dynamics across an entire gene body from nascent RNA sequencing data, we discovered that genes have two types of conserved regions near the disassociation site: one characterized by a T-rich region upstream of disassociation, and another characterized by a GC-rich region surrounding disassociation. Strikingly, the GC-rich disassociation regions have more accessible chromatin and higher levels of phospho-threonine 4 on the CTD of Pol II. Additionally, we find that upstream T-rich genes are preferentially affected by perturbations that alter disassociation, including heat-shock, viral infection, kinase inhibition, and arsenic treatment. Thus, our work has determined there are two types of Pol II disassociation regions, which are differentially affected by perturbation of cellular homeostasis.

## Introduction

The enzyme RNA Polymerase II (Pol II) is essential for life, as it transcribes all protein-coding genes, as well as many non-protein-coding and enhancer transcripts. The classical view of Pol II-mediated transcription describes three distinct phases: initiation, elongation, and termination. Moreover, Pol II transcription termination at genes consists of two steps: cleavage and polyadenylation of the mRNA and subsequent disassociation of Pol II from the DNA [1]. Although Pol II initiation and elongation are recognized as highly regulated steps that control gene expression, termination is often overlooked and assumed not to be a prominent driver of gene regulation. Additionally, research on Pol II termination to date has primarily focused on the first stage of termination, cleavage and polyadenylation, leaving the second part of Pol II termination, disassociation, understudied. Despite the lack of research on this process, transcriptional termination plays an important role in proper mRNA maturation, as altered termination can cause improper mRNA processing, disrupt mRNA export, and, in some cases, cause cell death [2–4]. Furthermore, certain stressors alter Pol II termination in a process known as run-on transcription, in which Pol II disassociates further downstream than normal [5, 6]. Run-on induced by heat-shock, oxidative stress, viral infection, and hypoxia results from altered RNA cleavage at a subset of genes [7–11]. However, whether stress-induced 3*′* alterations result solely from cleavage failure or also involve cleavage-independent mechanisms remains unclear. Therefore, it is evident that questions remain about the mechanistic details of Pol II termination under both control and perturbed conditions [12–14].

As Pol II transcription nears the end of a gene, mRNA is cleaved at the transcription cleavage site (TCS) and is subsequently released from polymerase. Notably, after pre-mRNA cleavage, Pol II continues to transcribe past the TCS along the DNA for some distance before disassociating from the DNA [15, 16]. In the prevailing model of Pol II transcription termination, known as the ”torpedo model” [1, 17–19], the nuclear 5*^′^*-3*^′^* exoribonuclease XRN2 is recruited to the uncapped 5*′* end of the nascent transcript and progressively degrades the RNA until it reaches Pol II, physically impacting Pol II processivity, and triggering disassociation.

Although the properties of the TCS are well-characterized for most transcripts, the locations of Pol II disassociation sites and any associated transcriptomic or genomic patterns remain unknown. Identifying Pol II disassociation sites is challenging, as nascent transcripts generated by Pol II after the TCS are rapidly degraded, making them difficult to capture in most biochemical assays. Moreover, experiments that directly capture Pol II positions, such as Pol II ChIP-seq, lack strand-specific information and suffer from high signal-to-noise ratios. In contrast, nascent run-on sequencing is a technique that captures transcripts actively being transcribed by any of the RNA polymerases [20, 21]. Unlike other assays, nascent run-on sequencing provides a dynamic snapshot of ongoing transcriptional activity, standing out as one of the few assays capable of capturing Pol II activity beyond the TCS.

Despite many efforts aimed at identifying the end of each transcript [22–25], the disassociation position remained difficult to reproducibly infer from any sequencing data [25]. This is mainly because most studies infer Pol II disassociation sites by identifying the most 3*^′^* position of read coverage, a position heavily dependent on data depth and quality. To address this issue, we developed a novel Pol II activity model that explicitly models the position of Pol II slowing that precedes disassociation [26, 27]. The slowing of Pol II induces a peak in nascent run-on data, which we refer to as the disassociation peak. By focusing on this peak, our approach is less susceptible to data depth and quality issues that influence the precise positioning of the end of transcription.

Here, we leverage our recently published Pol II model, a probabilistic mixture model that captures the four distinct stages of Pol II activity, namely loading, initiation, <u>e</u>longation and termination stages (Figure 1A) [26], on a variety of cell types in both basal and perturbed conditions [28]. We identify two broad classes of genes with distinct Pol II activity patterns regarding transcription termination. These two classes of genes have distinct chromatin accessibility profiles, as well as different levels of Pol II CTD threonine 4 phosphorylation (Th4P) near their sites of disassociation. These two classes have two evolutionarily conserved base composition patterns near the disassociation peak. Further, we find that the GC-rich regions near disassociation peaks reduce Pol II run-on transcription under stress. Our work provides insight and raises new questions regarding the mechanisms of Poll II disassociation and its role in affecting cell viability across conditions.

**Fig. 1.**
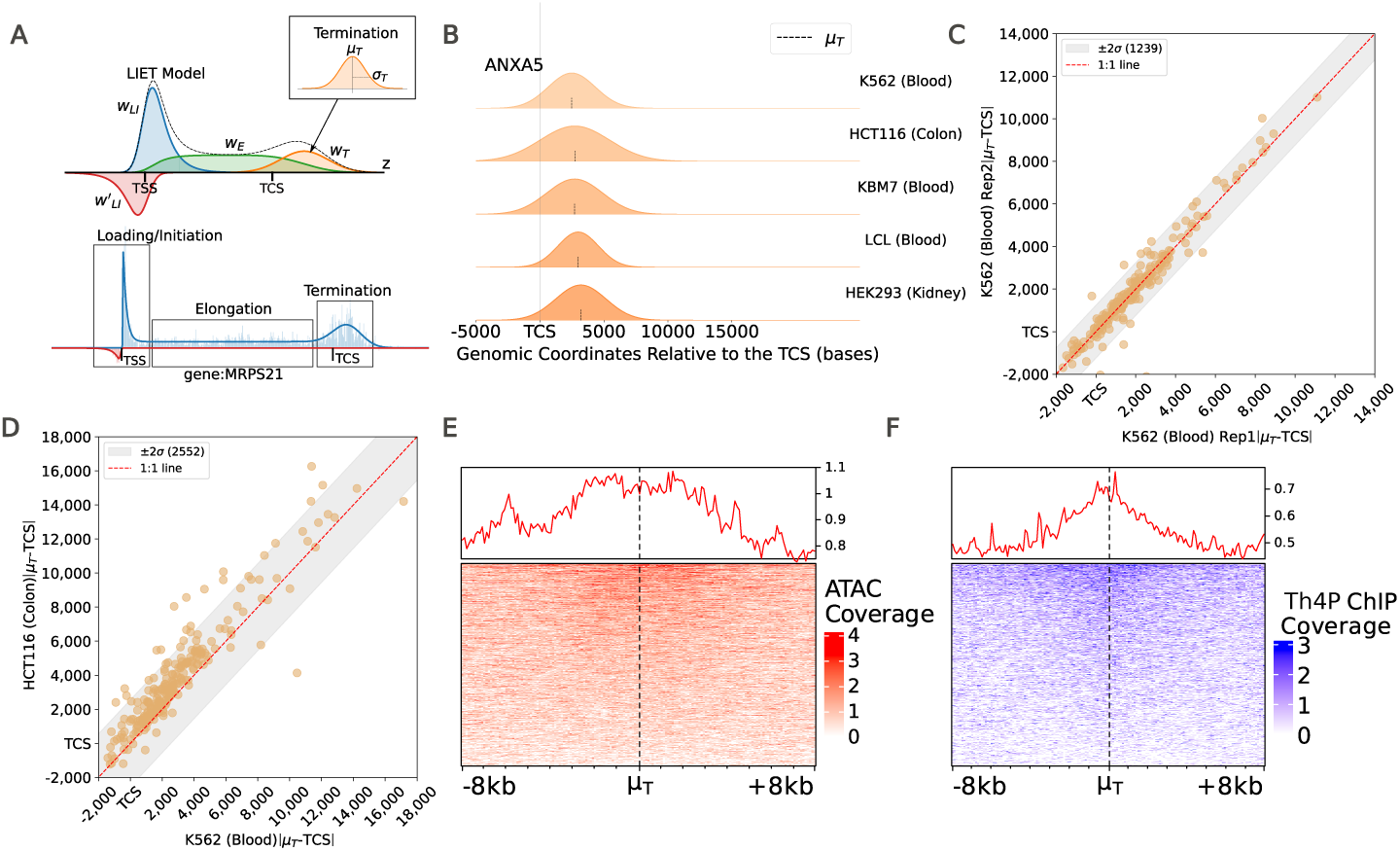
Disassociation positions are reproducible across human cell types and have elevated levels of Pol II CTD threonine 4 phosphorylation (Th4P) and chromatin accessibility. **A**. The LIET model parametrizes transcription on a per-gene basis. Termination is modeled as a Normal distribution, where *μ_T_* is the location of the disassociation peak and σ*_T_* captures variance (i.e., width of peak). **B**. LIET modeled disassociation peak (located at *μ_T_*) for the gene ANXA5 across human K562, HCT116, KBM7, LCL, and HEK293 cells (meta-sample for each cell type, Supplemental Table 1) relative to the TCS. **C**. |*μ_T_* − *TCS*| between replicates in human K562 cells (basal conditions). Grey region indicates technical variation (2*σ* = 1239bp) from the identity line. **D**. |*μ_T_* − *TCS*| between two different meta-samples of different cell types (HCT116 vs. K562). Grey zone is 2*σ* from the identity line (2552bp). **E**. ATAC-seq coverage at *μ_T_* in human lymphoblastoid cells. **F**. Phospho-mark threonine-4 (Th4P) ChIP-seq coverage at *μ_T_* in human HEK293 cells [29].

## Results

### The position of disassociation is reproducible across replicates and cell types

We sought to utilize the LIET model [26] across a broad set of nascent run-on data [28] to gain insight into the mechanisms of Pol II disassociation. Our model (LIET) captures the stages of transcription in the aggregate signals induced by Pol II (each represented by a single read) active within the bulk sample (i.e., the shapes of the data, Figure 1A). LIET fits directly to data, without considering sequence information, and has high accuracy and precision of parameter estimates across a broad range of noise levels, nascent run-on protocols, and read-depths. In the original LIET paper, we focused primarily on validating the model [26]. Thus, for that paper we ran LIET on only 163 manually curated, transcriptionally isolated genes transcribed in 24 cell types (to avoid confounding signals from overlapping transcription). While this analysis validated our mathematical model and served as a great starting point for a broader study on Pol II termination, the limited number of genes restricted our ability to determine what other features correlate with the position of disassociation.

Consequently, to study the features associated with disassociation, our first step was to expand our analysis to a larger set of transcriptionally isolated genes. To this end, we eliminated the requirement that the gene must be transcribed in all available cell types, thereby allowing us to expand the number of genes analyzed per cell type. We combined high-quality replicates for each cell type to create meta-samples with maximized depth and complexity, enabling more genes to achieve sufficient coverage for LIET to annotate. For every meta sample (per cell type), we identified a set of reasonably transcribed genes (depth cutoff: *≥* 1 read per kb) that contained no nearby gene annotations as well as no bidirectional transcription for the given cell type near Pol II disassociation (see Methods for full description). Utilizing these strategies, we expand the gene list to 1000-1500 genes per cell type, increasing the likelihood of identifying disassociation-associated features. As the gene lists are not perfectly overlapping across cell types, we fit LIET to a total of 3923 genes.

Applying LIET to this expanded gene set, we first sought to recapitulate our previous findings [26], namely that the position of disassociation was reproducible across cell types under basal and unperturbed conditions. The LIET model captures the disassociation peak in nascent sequencing data (the peak reflective of Pol II slowing prior to disassociation) as a Normal distribution N(*µ_T_*,*σ_T_*). Conceptually, the center of the Normal distribution (*µ_T_*) reflects the position most polymerases have reached prior to disengaging from the DNA. In contrast, the width of the peak (*σ_T_*) captures the biological and technical variation of the disassociation position. Importantly, we assume that Pol II disassociation occurs immediately after Pol II slowing.

Leveraging LIET, we found that the position of the Pol II disassociation peak (*µ_T_*) was reproducible across replicates and cell types in basal conditions for the larger set of genes (Figure 1A-D) [26]. In particular, we note that the variability of *µ_T_* was minimal when comparing matched samples (same protocol and laboratory of origin and with similar read depths) (Figure 1C). However, we observed that variability in the disassociation position increased when comparing samples prepared by different groups and at varying read depths (Figure 1D). To quantify this technical variation, we assumed measurement noise was randomly distributed around the expected identity line (1:1 relationship) between replicates and then modeled this variation as a Normal distribution. We subsequently used a 95% confidence interval (2\**σ*) to establish the expected range of measurement variation.

All pairwise comparisons between human cell types exhibited a linear relationship in *|µ_T_ − TCS|* values (average *r*^2^ = 0.8), with only 6.6% of genes deviating beyond *±*2*σ* from the identity line on average (Supplemental Figure S1). For the genes with *|µ_T_ − TCS|* values that fell outside of the 95% confidence interval, we found the variation in *µ_T_* was largely due to differences in *σ_T_* values between genes. For a gene with a larger *σ_T_* value, *µ_T_* became more variable (Supplemental Figures S2,S3A-F), consistent with its lower fidelity. These results reaffirm that the position of the Pol II disassociation peak across the genes we surveyed is consistent across human cell types and replicates.

Our extended gene list and newly annotated *µ_T_* values enabled us to investigate cellular features that correlate with the position of disassociation. We first wondered whether regions around the disassociation peak (measured by LIET using *µ_T_*values from PRO-seq data generated from the same lymphoblastoid cells) exhibited accessible chromatin. To address this, we generated and analyzed chromatin accessibility data from human lymphoblastoid cells (see Methods). Our analysis revealed that chromatin accessibility increased near the disassociation peak relative to the TCS (Figure 1E, Supplemental Figure S4). Overall, while it remains unclear if chromatin accessibility directly influences disassociation, our analysis indicates that chromatin becomes more accessible near the disassociation peak.

Additionally, phosphorylation on Pol II’s CTD is a prominent regulator of gene expression, aiding Pol II in recruiting regulatory factors during transcription. Th4P was recently reported to be enriched near sites of Pol II disassociation [29, 30]. Thus, we next examined Th4P ChIP-seq data [29] in comparison to the annotated *µ_T_* values from PRO-seq data in HEK293 cells (matching cell type). We confirmed that there are elevated levels of Th4P at *µ_T_* relative to the TCS (Figure 1F, Supplemental Figure S5). Although the exact role of Th4P in termination remains unclear, we hypothesize that this phospho-mark plays a role in regulating Pol II disassociation.

### GC content impacts Pol II disassociation position

The strong reproducibility of the position of the disassociation peak prompted us to hypothesize that the nucleotide content downstream of the cleavage site may play a role in the location of disassociation. To investigate this, we examined the base composition near *µ_T_*across all genes and observed increased T-content upstream of *µ_T_* . We defined this pattern as an “upstream T-rich” pattern (Supplemental Figure S6).

To determine whether this base composition pattern was consistent across all genes analyzed, we clustered our genes based on the AT% surrounding the site of Pol II disassociation (*±* 3kb). Using K-means clustering with different numbers of clusters (k=2-5), we consistently identified two patterns, regardless of the number of clusters selected: a small subset (*≈*12% of all genes analyzed) with a GC-rich pattern at *µ_T_* and a larger subset with a T-rich pattern upstream of *µ_T_* . With increasing numbers of clusters (*k >* 1), the upstream T-rich pattern would split into subsets that varied by where the T-enrichment ended (Supplemental Figure S7A-D). Next, we sought to determine whether these two patterns were discrete or part of a continuous spectrum. To answer this question, we ranked genes based on their GC% (*±* 500pb) around *µ_T_* and examined quantiles, looking for the inflection point between the GC-rich and the upstream T-rich gene sets (Figure 2A, Supplemental Figure S8A). From this, we define two separate gene groups with different base composition patterns near disassociation, one smaller subset rich in GC-content and another larger subset with the upstream T-rich pattern. While we define these as separate groups, the distinction between them exists along a spectrum (Supplemental Figure S8A).

**Fig. 2.**
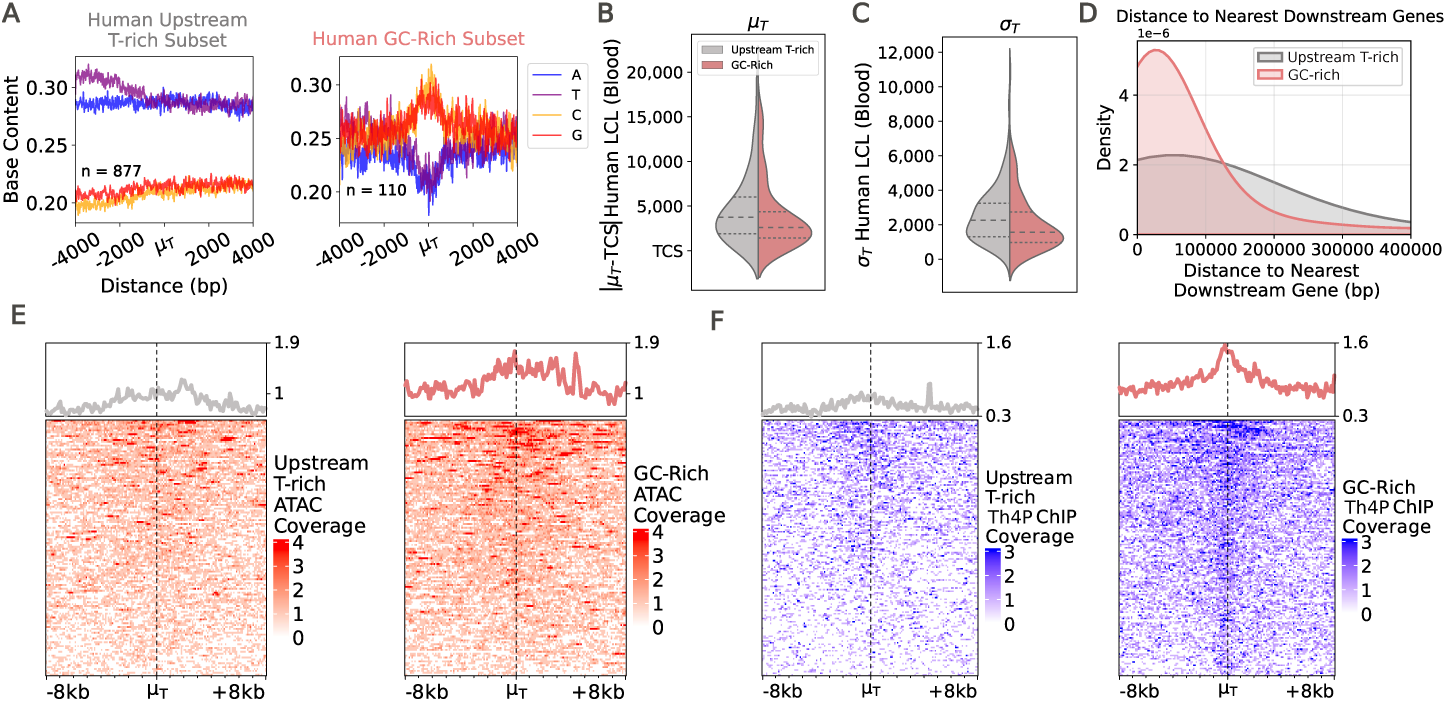
Two subsets of genes with distinct base composition patterns and disassociation properties. **A.** Two base composition patterns were identified. The dominant pattern occurs in 877 of surveyed genes and is rich in T content upstream of the disassociation peak. The smaller pattern is GC-rich at the disassociation peak (*μ_T_*). Background and TCS base composition are shown in Supplemental Figures S20-S21. **B**. Violin plots of the distance traveled after TCS (|*μ*_T_ − TCS|) and **C**. *σ_T_* . Upstream T-rich genes were randomly sub-sampled to equal the same number of GC-rich genes (n=110). **D**. Density plot of the distance from the 3′ end of the 3′ UTR to the 5′ UTR of the nearest downstream MANE transcript [62] at GC-rich genes (pink) vs. upstream T-rich genes (grey). All genes called in at least one sample utilized, with the T-rich set sub-sampled to match the GC-rich set size (n=275). **E**. Chromatin accessibility (human lymphoblastoid cells, Supplemental Table 1) near disassociation peak (centered on *μ_T_*) at upstream T-rich genes (left) versus GC-rich genes (right). Upstream T-rich genes are randomly sub-sampled to equal the number of GC-rich genes (n=134). **F**. Th4P ChIP-seq coverage (human HEK293 cells, [29]) at upstream T-rich (left) and GC-rich (right) genes, Upstream T-rich genes are randomly sub-sampled to equal the number of GC-rich genes (n=159).

Interestingly, the two subsets of genes had distinct disassociation properties. We compared the two subsets by first randomly subsampling the upstream T-rich subset to match the number of genes in the GC-rich, controlling for differences in the number of genes between the two groups. Using these equally sized subsets, we find that Pol II disassociated closer to the TCS and showed greater precision in the disassociation peak position (i.e., smaller *µ_T_* and *σ_T_*values on average, Figure 2B-C) in the GC-rich gene subset compared to the upstream T-rich subset. Genes in the GC-rich subset also tend to lie in more gene-dense regions, with the average distance to the nearest downstream gene being smaller compared to the upstream T-rich gene set (Figure 2D). Notably, the GC-rich set of genes also had dramatically stronger Th4P signal [29] near *µ_T_* (Figure 2E). These regions were also generally more accessible (Figure 2F). These findings reveal two subsets of genes with distinct properties of disassociation, indicating that Pol II disassociation varies in a gene-specific manner.

### Disassociation-associated base composition patterns are evolutionarily conserved

We next wondered whether the two dominant sequence patterns observed at disassociation, GC-rich and upstream T-rich, were evolutionarily conserved. We hypothesized that if base composition near *µ_T_* was evolutionarily conserved, the nucleotide content likely impacts Pol II disassociation efficiency. To test this, we first examined a deeply sequenced, high-quality mouse PRO-seq dataset [31]. Using the same techniques that were used in human (clustering followed by quantile analysis), we find the same two gene sets, a GC-rich and an upstream T-rich subset, in mouse as was observed in human (Supplemental Figure S9A). The distance traveled after the TCS varied tremendously between the two species (Supplemental Figure S9B). However, a more detailed examination of these differences was stymied by the fact that the human and mouse samples were not matched (i.e., different cell types and non-overlapping gene sets), and that sequence alignments in regions outside genes are generally challenging [32, 33].

To address these issues, we conducted PRO-seq on lymphoblastoid cell lines from rhesus, gibbon, and squirrel monkey (see Methods), enabling direct comparisons with human lymphoblastoid samples. Data for each species were assessed for quality and mapped to their native genome. We identified shared orthologs between human and each target species using Benchmarking Universal Single-Copy Orthologs (BUSCO)[34], and obtained annotations for conserved orthologs using the Progressive Cactus multiple-genome aligner [35, 36]. Using these orthologs, we applied the same pipeline for isolated gene curation–meta-sample generation and coverage/bidirectional filtering–but omitted the annotation isolation filter to avoid excessively restricting the number of genes we analyzed. After curating the set of isolated genes in each species, we ran LIET on each species’ resulting gene set.

Using k-means clustering followed by quantile analysis on the base content near *µ_T_*, we found that both of the base composition patterns at *µ_T_* were recovered across the three non-human species (Figure 3A, Supplemental Figures S8B-D,S10A-B,S11A-C). Like human, Pol II disassociation was more precise and occurred closer to the TCS (smaller *µ_T_* and *σ_T_* values) in the GC-rich subset compared to the upstream T-rich subset in each species (Figure 3B-C, Supplemental Figure S12A-D).

**Fig. 3.**
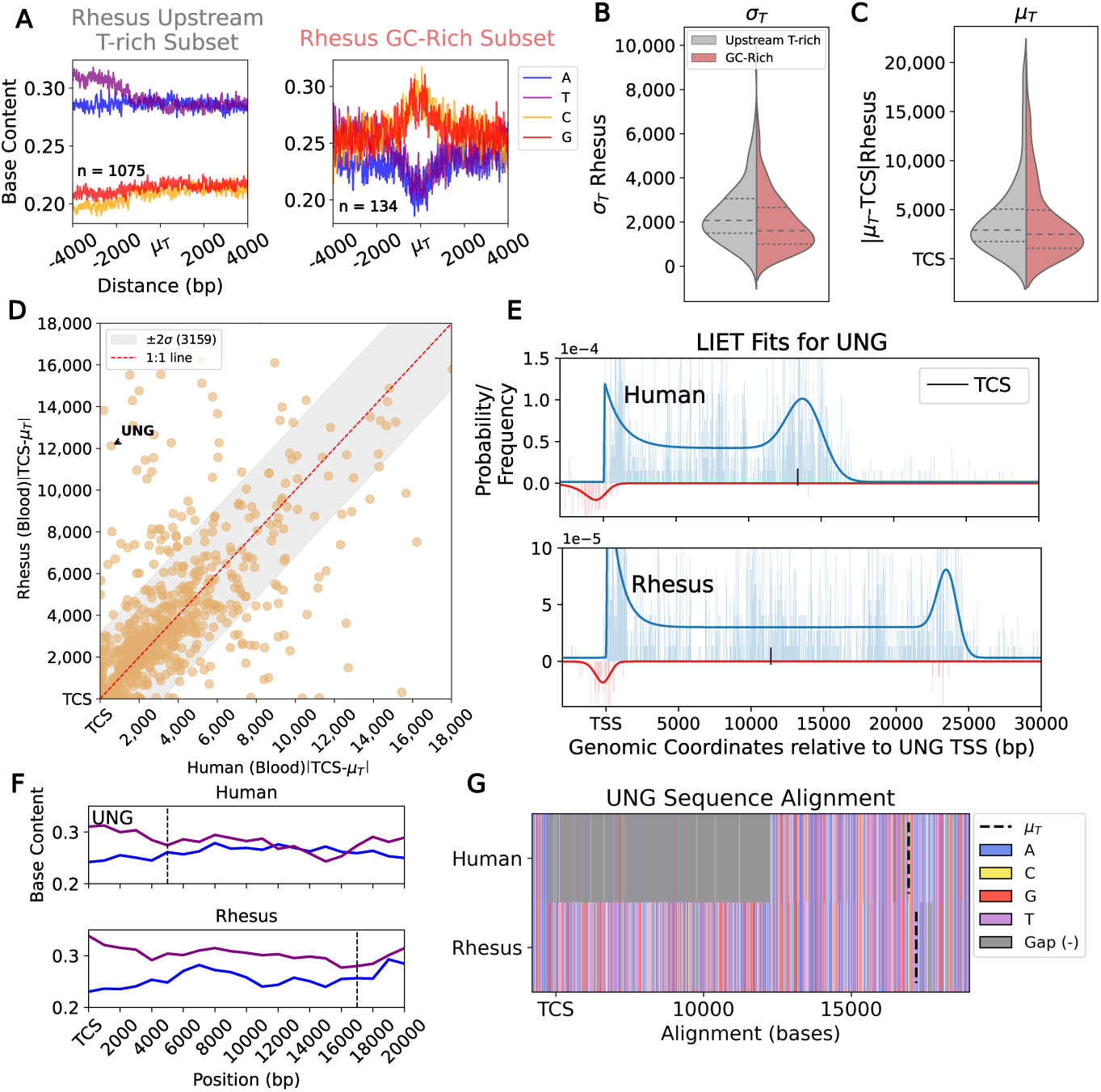
The GC-rich and upstream T-rich base composition patterns are evolutionarily conserved. **A**. Upstream T-rich and GC-rich base composition patterns near *μ_T_* in rhesus monkey (rheMac10 genome). Violin plot of **B**. *σ_T_* and **C**. the distance from the TCS to *μ_T_* in each subset of genes. Upstream T-rich genes were randomly sub-sampled to equal the same number of GC-rich genes (n=134). **D**. |*μ_T_* − *TCS*| in rhesus monkey vs human lymphoblast cells (combined replicates, meta-samples). Grey region calculated on replicates per species, showing rhesus sample which had larger variation. **E**. LIET fits for gene UNG in both human (above) and rhesus monkey (below). **F**. Base composition of A (blue) and T (purple) in region from the transcription cleavage site (TCS) to *μ_T_* (dashed line) in human and rhesus monkey for gene UNG. G. Gene UNG sequence alignment from the TCS to *μ_T_* shows an indel (grey in human) downstream of the cleavage site that shifts *μ_T_* upstream in human relative to rhesus.

Given the strong overall conservation of the base composition patterns near *µ_T_* across the species, we next sought to explicitly compare *µ_T_* between species. Remarkably, we found the distance traveled after cleavage (*|µ_T_ −TCS|*) is not well conserved (Figure 3D). In rhesus monkey, for example, the UNG gene disassociation peak is shifted further downstream relative to its position in human cells (Figure 3E). Additionally, looking at the base composition downstream of the TCS between the human and rhesus monkey’s genome, we observe that the T-rich region upstream of *µ_T_* is also shifted downstream in rhesus monkey at UNG (Figure 3F). This shift in base composition is reflective of the large indel downstream of the TCS (Figure 3G), which has led to a shortening of the length of Pol II processivity after cleavage in human relative to rhesus. Hence, the base composition patterns around the disassociation peak are well conserved despite shifts in the distance Pol II travels after the TCS across the species. These findings suggest that base composition patterns may drive the position of Pol II disassociation.

### Different perturbations influence the 3*^′^* end differently

Upon certain stressors, such as heat-shock (Figure 4A) and viral infections (Figure 4B), Pol II displays “run-on” behavior, leading to a downstream shift in the position of disassociation relative to normal conditions [5–7]. With this in mind, we next wondered whether these base composition patterns were also present at the downstream disassociation position associated with stress-induced run-on transcription. In our development of the LIET algorithm, we showed that our model can capture the new, downstream position of RNA Pol II disassociation in heat-shock conditions for the small original gene set [26]. Here, we first sought to expand our analysis of stress-induced run-on, both by examining more genes and considering additional perturbations available in the nascent database [28].

**Fig. 4.**
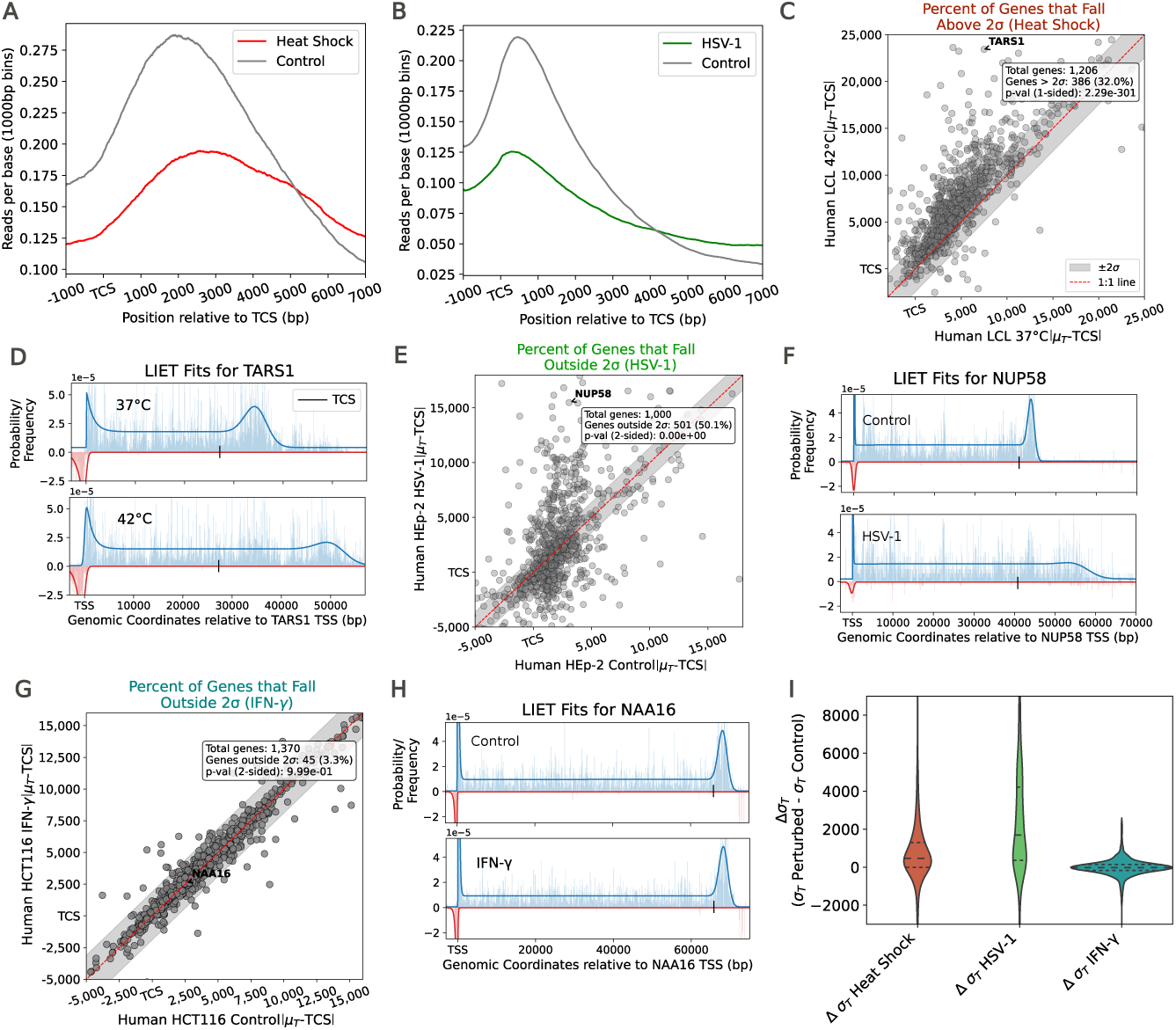
Each stressor impacts disassociation differently. LIET is run-on combined replicates and 2σ calculated from replicates in control conditions. **A.** Meta-gene plot of PRO-seq coverage in heat-shock downstream of the TCS in control (grey) versus heat-shock (red) conditions (human lymphoblastoid cells, [37]). **B**. Meta-gene plot of PRO-seq coverage in HSV-1 infection downstream of the TCS in control (grey) versus HSV-1 (green) conditions (human HEp-2 cells, [42]). **C**. |*μ_T_* −TCS| in control (x-axis) vs heat-shock (y-axis) conditions. One-sided binomial test finds significant run-on (p-value 2.29e-301) with 32% of genes shifting *μ_T_* downstream. **D**. LIET fits for gene TARS1 in control (37◦ C, top) and heat-shock (42◦ C, bottom) conditions. *μ_T_*. |*μ_T_* −*TCS*| in control (x-axis) vs HSV-1 (y-axis) conditions shows significant changes in disassociation position (two-sided binomial test pvalue ≈ 0.00). See Supplemental Figure S14 for 1-sided binomial tests on run-on and shortening. **F**. LIET fits for gene NUP58 in control (top) and HSV-1 (bottom) infection conditions. **G**. |*μ_T_* −*TCS*| in control (x-axis) vs IFN-γ (y-axis) conditions. IFN-γ does not impact the disassociation position of Pol II (2-sided binomial p-value = 9.99e-01) [41]. H. LIET fits for gene NAA16 in control (top) and IFN-γ (bottom) conditions. I. Change in σT between perturbed and control conditions (Δσ*_T_*).

First, we ran LIET on our expanded gene list on nascent RNA-sequencing data obtained from cells subjected to heat-shock [37]. In this experiment, human lymphoblastoid cells were exposed to 42*^◦^*C for 1 hour before performing PRO-seq. After running LIET on these samples, we observed a downstream shift in disassociation across many isolated genes (n=1206 genes; Figure 4C). Notably, the magnitude of the stress-induced shift varied across genes, with some genes exhibiting minimal changes to *µ_T_* (within the range of technical variation), while others exhibited shifts that exceeded these bounds. To formalize a quantitative definition of run-on transcription, we define run-on as occurring when a gene’s measured *µ_T_* under stress shifts beyond the measurement error bounds (95% confidence interval) obtained in the control replicates. Furthermore, we can assess for stress-induced run-on (or shortening) with a one-sided binomial test and for general alterations in the disassociation peak (changes in either direction) with a two-sided binomial test. We applied this run-on classification approach to the heat-shock data, where we identified a statistically significant change in the position of disassociation (p-value = 2.29e-301) at many genes (n=386), such as TARS1 (Figure 4D).

We next asked whether there was a base composition pattern at the run-on associated *µ_T_* values in heat-shock. Under stress, Pol II bypasses canonical disassociation sites at certain genes–potentially due to alterations in Pol II kinetics and/or cleavage failure [5, 8]. We hypothesized that when disassociation is perturbed, Pol II may encounter an alternative base composition pattern that assists in disassociation at the new position. To test this, we looked for base-composition patterns near *µ_T_* in the perturbed conditions. We found no discernible base composition properties near *µ_T_* under perturbed conditions (Supplemental Figure S13A), suggesting that Pol II disassociation may occur through a base composition-independent mechanism under stress.

Next, we sought to apply our run-on classifier to another commonly reported condition that induces transcriptional run-on, viral infection [5–7]. HSV-1 alters transcription termination by interfering with the cleavage and polyadenylation factor machinery [5, 10, 38, 39]. Birkenheuer et al. infected aryngeal epidermoid carcinoma cells with HSV-1 and ran PRO-seq [40]. We first used the standard meta-gene approach to confirm the presence of run-on in this dataset upon exposure to HSV-1 infection (Figure 4B). Consistent with the meta-gene, the LIET analysis approach finds that HSV-1 induces a downstream shift in disassociation at a subset of genes (n=269, p-value=2.33e-188, Supplemental Figure S14A, Figures 4E-F). Surprisingly, we also found that HSV-1 induces an upstream shift at a separate, slightly smaller group of genes (n=232, p-value=4.37e-147, Supplemental Figure S14B). Notably, the metagene approach (Figure 4B) masks gene-specific shortening of *µ_T_* by averaging across many genes. Additionally, like heat-shock, we found no clear base composition properties near *µ_T_* under HSV-1 infection (Supplemental Figure S13B-C). LIET’s capability of assessing stress-induced changes to disassociation on a per-gene level clearly identifies distinct gene subsets exhibiting altered run-on behavior in response to HSV-1 infection.

Additionally, we wondered whether the HSV-1-induced changes in disassociation were due to the host interferon response or the HSV-1 virus itself [39]. We examined IFN-*γ* exposure in lymphoblastoid cells [41] and found that it did not impact how far Pol II traveled before disassociating (Figure 4G), as shown at the gene NAA16 (Figure 4H) in this cell type. This result implies that HSV-1-induced changes in disassociation are likely due to the pathogen rather than the host response ([42]). Furthermore, these findings demonstrate that alterations to *µ_T_* under stress are stimuli-specific.

Finally, we noticed that the efficiency of Pol II disassociation (width of disassociation peak–*σ_T_*) differed substantially between heat-shock and HSV-1 infection. *σ_T_* was large in HSV-1 compared to either heat-shock or IFN*γ* (Figure 4I), indicating that the disassociation peaks became less defined and disassociation positions were overall less precise in infection. We see this trend reflected in the gene NUP58, where the disassociation peak widens and appears more diffuse under infection, relative to control conditions (Figure 4F).

### Under stress, Pol II runs-on less frequently at GC-rich genes

Next, we asked whether the base composition characteristics of standard (control condition) transcription (upstream T-rich or GC-rich at disassociation) affect whether the gene exhibits run-on transcription. We address this by investigating whether genes with run-on were enriched in the GC-rich or upstream T-rich subsets. As expected, we found no enrichment for either subset in the IFN-*γ* data, consistent with the lack of run-on in this condition (Figure 5A). Interestingly, we found that in both heat-shock (Figure 5B) and HSV-1 (Figure 5C), the run-on set of genes was enriched for only the upstream T-rich set (p-value *<* 0.05). This finding suggested that the GC-rich subset of genes was less prone to stress-induced run-on.

**Fig. 5.**
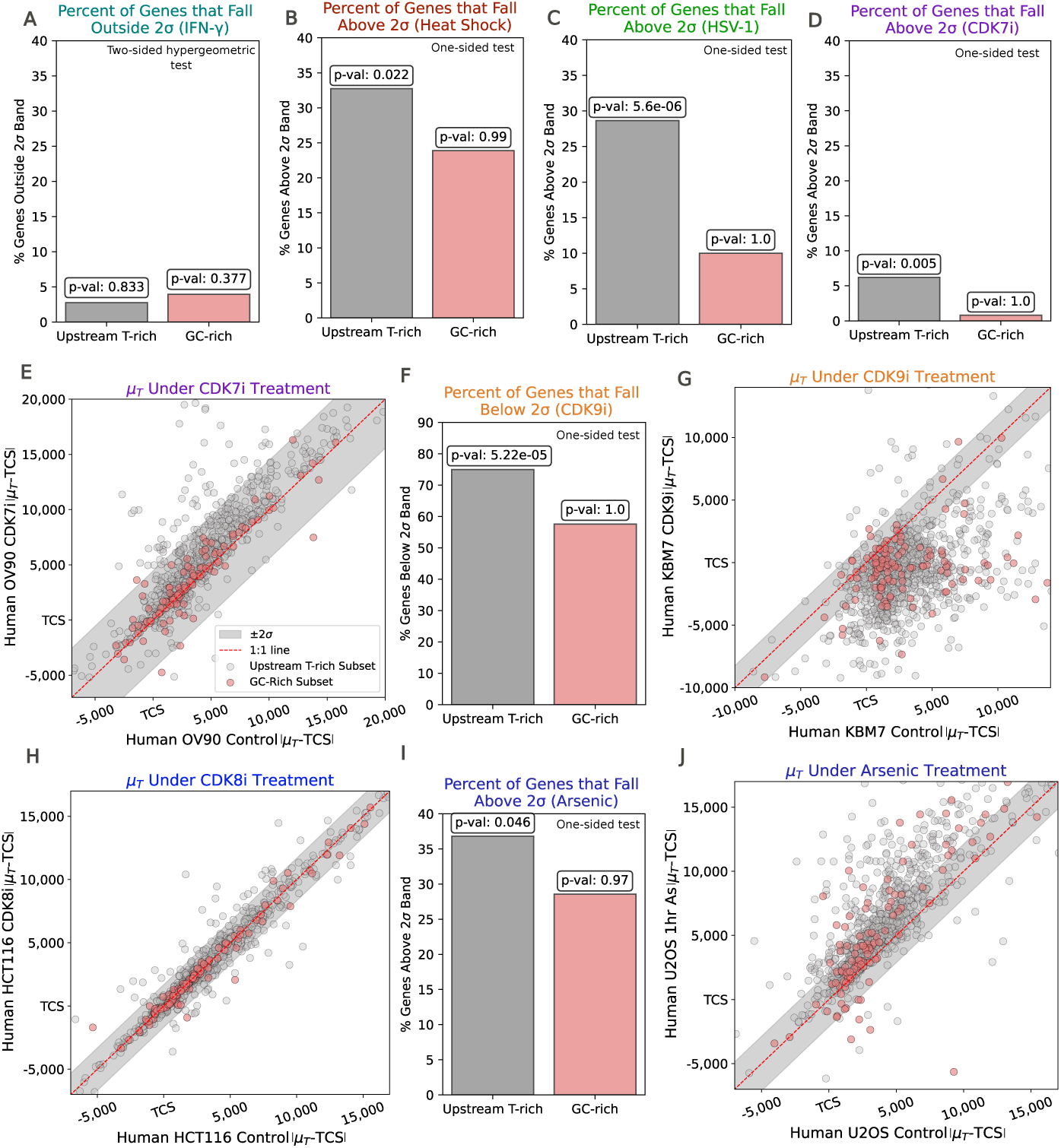
GC-rich genes are less prone to stress induced run-on. A hypergeometric test for enrichment of genes from the upstream T-rich subset (grey) and GC-rich subset (pink) for **A**. IFN-γ- induced changes in disassociation [41], **B**. heat-shock-induced run-on [37], **C**. HSV-1 infection-induced run-on [40], and **D**. CDK7 inhibition-induced run-on [64]. **E**. |*μ_T_* − TCS| in control (x-axis) versus CDK7 inhibition (y-axis) [64]. Scatter points are colored on upstream T-rich (grey) versus GC-rich (pink) subsets. **F**. Hypergeometric test on CDK9 inhibition experiment for genes that shorten [45]. G. |*μ_T_* − *TCS*| in control (x-axis) versus CDK9 inhibition (y-axis) conditions [45]. Scatter points are colored on upstream T-rich versus GC-rich subsets. **H**. |*μ_T_* −*TCS*| for control (x-axis) vs CDK8 inhibition (y-axis) conditions (HCT116 CDK8 inhibition analog sensitive cell line) [41]. Scatter points are colored on upstream T-rich versus GC-rich subsets. **I**. Hypergeometric test on arsenic experiment for run-on. J. |*μ_T_* −*TCS*| for control (x-axis) vs arsenic treatment (y-axis) conditions. Scatter points are colored on upstream T-rich versus GC-rich subsets.

Given this result, we next tested whether the subset of genes with T-rich typical disassociation sites was also enriched among run-on genes in other conditions. As a first pass, we turned to our previously published CDK7 inhibition (by SY5609 in human OV90 cells) experiments, where our prior work had found run-on transcription by a meta-gene analysis[43, 44]. LIET revealed that CDK7 inhibition shifted *µ_T_* downstream in a smaller subset of genes (n=115, Supplemental Figure S14C) compared to heat-shock or HSV-1 infection. Nevertheless, upstream T-rich genes were enriched among run-on genes in this experiment (p-value 0.005, Figure 5D-E), consistent with heat-shock and HSV-1 infection. Additionally, as with heat-shock, the precision of the new disassociation position remained largely unchanged (Supplemental Figure S15).

Next, we wondered whether inhibiting other CDK proteins would similarly lead to run-on transcription. To address this, we separately examined both CDK9 inhibition (by NVP-2) [45] and CDK8 inhibition (in a CDK8 analog-sensitive HCT116 cell line) [41]. CDK9 and CDK8 each regulate distinct regulatory events at gene 5*′*-ends [46–48]. CDK9 inhibition induced a global upstream shift in disassociation, suggesting extensive (n=840, Supplemental Figure S14D) shortening in the extent of transcription (Figure 5F-G), consistent with previous reports of premature termination [49]. Interestingly, CDK9 inhibition also showed reduced precision in Pol II disassociation upon drug treatment, a similar but less pronounced trend to HSV-1 infection (Supplemental Figure S15). Intriguingly, previous reports also link CDK9 to XRN2 activity [50], which may be leading to the loss of precision. Additionally, upstream T-rich genes showed significant enrichment for upstream disassociation upon CDK9 inhibition (p-value = 5.22e-05), while GC-rich genes did not (Figure 5F-G). In contrast, CDK8 inhibition showed no signs of run-on transcription (2-sided p-value=0.69, Figure 5 H, Supplemental Figure S14E), similar to the IFN-*γ* data.

Lastly, we considered two environmental exposure datasets, wood smoke particles [51] and arsenic exposure. Similar to CDK8 inhibition and IFN-*γ*, wood smoke did not lead to changes in the position of disassociation (Supplemental Figure S14F). However, arsenic exposure (in human U-2 OS cells, prepped in-house, see Methods), which induces oxidative stress, resulted in a downstream shift in disassociation positions (n=289, Supplemental Figure S14G) reminiscent of heat-shock. The arsenic exposure samples had lower sequencing depth and complexity than other samples, resulting in high replicate variation in *|µ_T_ −TCS|* among the control samples (Supplemental Figure S16A). To ensure quality fits despite the lower complexity, we required higher coverage downstream of the TCS (see Methods), which reduced technical variation between control replicates (Supplemental Figure S16B). With this adjustment, the hypergeometric test showed that the upstream T-rich subset was enriched among run-on genes (p-value 0.046; Figure 5I-J), consistent with other run-on-inducing conditions (heatshock, HSV-1 infection, and CDK7 inhibition). Notably, without the adjusted depth and complexity filter, the trend remained directionally consistent but was not statistically significant. These findings suggest that GC content near *µ_T_* appears to be an important regulator in the fine-tuning of where Pol II disengages under perturbation, as higher GC content near *µ_T_* leads to fewer 3*′* end alterations.

## Discussion

Termination is the least well-understood phase of transcription. While the first stages (loading/initiation and elongation) are highly regulated modulators of gene expression, it remains unclear whether similar regulatory mechanisms are in place at the 3*′* end of transcription. Using the LIET model, we identified positions of Pol II disassociation in PRO-seq data across 9 cell types, 5 species, and 7 different perturbations to gain insights into the mechanism of transcription termination under control and stressed conditions.

Our analysis revealed two classes of genes, each defined by distinct, evolutionarily conserved base-composition patterns near *µ_T_* . The first class exhibited elevated GC content near the site of disassociation (*≈* 12% of the genes analyzed), while the second class displayed enrichment of T nucleotides upstream of the disassociation peak. While we distinguish these as separate groups to gain insight into the dynamics between base composition and disassociation, it’s important to note that sequences analyzed exist along a spectrum rather than fall into discrete groups. At the GC-rich genes, Pol II transcribed for shorter distances past the cleavage site before disengaging from the DNA, and disassociation occurred with higher positional precision (small *σ_T_*values). Additionally, we found elevated CTD Th4P phosphorylation (a termination-associated mark) and chromatin accessibility near disassociation peaks in these genes relative to the upstream T-rich gene set. Furthermore, perturbation-induced run-on was less prevalent in the GC-rich set of genes compared to the upstream T-rich genes (Figure 5, Supplemental Figures S17A-E,S18A-B). Elevated GC content at the 5*^′^* end of genes promotes Pol II promoter-proximal pausing prior to elongation [52]. Therefore, it is plausible that increased GC content at the 3*′* end of transcription may contribute to de-accelerating Pol II after cleavage, allowing more time for XRN2 to catch up and facilitate efficient termination.

We assessed disassociation only at transcriptionally isolated genes to avoid complications arising from overlapping transcription. In regions of overlapping transcription, such as intronic enhancer RNAs [28], the LIET model can produce poor fits due to ambiguity in read assignment. This occurs because it is hard to determine which reads originated from which transcript in overlapping regions. The results here (across all experiments) evaluated disassociation at 3923 genes. Due to this limitation, it remains to be seen whether additional base composition patterns around disassociation exist beyond the two identified here. Additionally, the GC-rich subset was relatively small (n = 275 genes across all experiments), limiting the ability to identify additional motifs or conduct pathway or gene set enrichment analyses. Importantly, comparisons between GC-rich and upstream T-rich genes were performed by randomly subsampling the T-rich group to match the GC-rich group size, controlling for differences in gene number (Figures 2,3, Supplemental Figure S17A-E). In the future, identifying disassociation positions genome-wide, including in transcription-dense regions, will enable more rigorous pathway analyses and reveal whether additional base composition patterns exist beyond the two identified here.

Most studies to date have assessed Pol II disassociation from a meta-gene perspective, averaging all signals downstream of the cleavage site across genes. While this method helps identify global trends, particularly in run-on conditions [5, 44, 53], it struggles to detect distinct patterns within subsets of genes. In this study, we took a different approach, using the LIET model to identify disassociation peak sites on a per-gene basis and intersecting those positions across a variety of omics datasets to gain insight into the mechanism(s) of disassociation. By applying the LIET model to evaluate perturbation-induced changes in disassociation on a per-gene basis, we uncovered distinct run-on patterns that vary by stressor. Heat-shock, HSV-1 infection, CDK7 inhibition (SY5609), and arsenic exposure all triggered run-on transcription, yet the number of affected genes and the distances traveled differed substantially across conditions.

Notably, we observed that disassociation not only shifts downstream under stress but also sometimes shifts upstream. For example, treatment of chronic myelogenous leukemia cells with the CDK9 inhibitor NVP-2 induced a global upstream shift in disassociation, while HSV-1 infection caused an upstream shift in a subset of genes. Additionally, NVP-2 and HSV-1 infection also reduced the precision of Pol II disassociation peak positions, widening the 3*′* slowing peak in nascent sequencing data. Though failed or altered cleavage often precedes run-on transcription [7], it remains unclear why Pol II runs-on to different extents across perturbations. Our findings suggest that both the magnitude and precision of these disassociation changes are largely stressor-specific.

Stress-induced alterations in transcription termination may have profound implications for cellular behavior, leading to altered mRNA production [2]. For example, when cleavage at the TCS fails, the transcript cannot be processed and exported to the cytoplasm for translation [2, 7]. Additionally, given that many human protein-coding genes are positioned in tandem [54, 55], alterations in Pol II disassociation can cause a phenomenon known as read-in transcription [56], where Pol II continues to transcribe beyond the cleavage site into the downstream gene. Read-in transcription can negatively impact the viability of the transcripts produced by the downstream gene [1]. Notably, stress-induced run-on was less prevalent in the GC-rich subset of genes compared to the upstream T-rich subset. Thus, elevated GC content could be an important feature in limiting run-on transcription at certain genes under stress. Since read-in of downstream genes can cause translational repression through intron retention [57], higher 3*′* GC content may serve as a protective mechanism to prevent inadvertent silencing of neighboring genes. This would be beneficial for protecting metabolic and developmental genes critical to cell survival, even under stress conditions that induce widespread transcriptional repression.

Overall, this analysis identified transcriptomic and genomic patterns associated with Pol II disassociation, including base composition, chromatin accessibility, and Pol II CTD Th4P levels. By investigating disassociation under control and perturbed conditions, we identified multiple gene classes with distinct disassociation properties. This work reveals that 3*′* end transcription regulation may be more complex than previously appreciated and provides a foundation for further investigation of transcription termination.

## Methods

### Publicly Available Data

Publicly available data were obtained from the DBNascent repository [28]. To ensure sufficient sequencing depth, we analyzed 8 human cell types from larynx, blood, colon, ovary, lung, kidney, and bone tissues, and 5 species across 2 cell types from blood and blastocyst tissues. Furthermore, we created meta-samples, combining reads from all technical and biological replicates for each cell type (basal analysis) and condition (perturbation analysis) using the same methods as previously published[26].

### Chromatin Accessibility Assay (ATAC-seq)

We obtained the lymphoblastoid cell line (ID: COMIRB 08–1276) from the Translational Nexus Biobank, University of Colorado School of Medicine, JFK Partners [37]. The cell line was derived from an individual’s blood sample and immortalized by infecting it with Epstein-Barr virus. Lymphoblastoid cultures were grown in RPMI media supplemented with 20% FBS, 100 U/ml penicillin, 100 *µ*g/ml streptomycin, and 2mM L-glutamine, and were maintained in incubators with 5% CO2 at 37*^◦^*C before harvesting. The samples were provided fully de-identified, and the authors had no access to identifying information about the individual.

### ATAC-seq

ATAC-seq library was made using the [58] protocol. In short, 50,000 viable lymphoblastoid cells were pelleted by centrifugation at 500 x g for 5 minutes at 4*^◦^*C. All supernatant was aspirated down to 100 *µ*l, avoiding the visible cell pellet. 50 *µ*l of resuspension buffer (water with 10 mM Tris-HCl pH 7.4, 10 mM NaCl, 3 mM MgCl_2_, 0.1% IGEPAL, 0.1% Tween-20, 0.01% Digitonin) and the cells re-suspended pipetting up and down 3 times, and incubated on ice for 3 minutes. Then, 1 *µ*L of re-suspension buffer (water with 10 mM Tris-HCl pH 7.4, 10 mM NaCl, 3 mM MgCl_2_, 0.1% Tween-20) was added and the tube was inverted 3 times to mix. Nuclei were then pelleted again by centrifugation at 500 x g for 10 min at 4*^◦^*C. The supernatant was then aspirated (avoiding the visible cell pellet). The pellet was then re-suspended by pipetting 6 times with 50 µL of the transposition mix (25 µL Tagment DNA Buffer Illumina Ref. 15027866, 2.5 µL Tagment DNA Enzyme 1 Illumina Ref. 15027865, 0.5 µL Digitonin diluted 1:1 with water, 0.5 µL 10% Tween-20, 5 µL water, 16.5 µL PBS), and were incubated for 30 minutes in a heat block at 37*^◦^*C. The sample was then cleaned using the DNA Clean and Concentrator-5 Kit (Zymo Research Ref. D4014) following the manufacturer’s instructions, and eluted in 21 µL elution buffer. Then, the library was amplified for 5 cycles with PCR using the NEBNext IIx Master Mix (NEB Ref. M0544S). Tubes were then removed from the thermocycler and stored on ice. Then, using 5 *µ*L (10%) of the pre-amplified mixture, a 15 *µ*L qPCR was run to determine the number of additional PCR cycles needed. The post-amplified libraries were then cleaned up with the DNA Clean and Concentrator-5 Kit (Zymo Research Ref. D4014) and eluted in 20 *µ*L of water.

### Cross Species Nascent Run-On Assay (PRO-seq)

#### Cell Conditions and Nuclei Isolation Across Species

Rhesus (ID: Mm 290-96, obtained from Yoav Gilad Lab), gibbon (ID: Ricky, obtained from Lucia Carbone Lab), squirrel monkey (ID: SML clone 4D8, obtained from ATCC Ref. CRL-2311) and human (ID: GM12878, obtained from Coriell/NIGMS) lymphoblastoid cells were treated with 0.1% BSA in PBS (final concentration of 0.00004% BSA) 1 hour prior to nuclei isolation. Nuclei isolation was performed as described in [59]. After treatment, lyphoblastoid cell cultures (10-30 million cells) were washed twice in ice-cold PBS. Next, cell pellets were re-suspended in 6 mL of lysis buffer (0.1% DEPC-DI water with 10 mM Tris-HCl pH 7.4, 2 mM MgCl_2_, 3 mM CaCl_2_, 0.5% IGEPAL, 10% Glycerol, 1 mM DTT, Invitrogen Ref. AM2696 SUPERase-IN RNAse inhibitor, and with Roche Ref. 11836170001 protease inhibitor cocktail) and centrifuged for 15 minures at 4*^◦^*C at 1000 x g. The pellets were then re-suspended in 1mL lysis buffer using Finntip wide orifice pipette tips (Thermo Scientific Ref. 9405163). The re-suspended pellets were then mixed with 4 mL more of lysis buffer, and centrifuged a second time for 15 minutes at 4*^◦^*C at 1000 x g. The pellets were then re-suspended a second time in 1 mL of lysis buffer (using Finntip wide orifice pipette tips), transferred to low binding 1.7 mL eppendorf tubes (Costar Ref. 3207), and centrifuged for 5 minutes at 4*^◦^*C at 1000 x g. The pellets were then carefully re-suspended a third time using Finntip wide orifice pipette tips in 500 µL freezing buffer (0.1% DEPC-DI water with 50 mM Tris-HCl pH 8.0, 5 mM MgCl_2_, 40% Glycerol, 0.1 mM EDTA pH 8.0, and SUPERase-IN RNAse inhibitor), and centrifuged for 2 minutes at 4*^◦^*C at 2000 x g. The resulting nuclei pellets were re-suspended a final time in 110 µL of freezing buffer using Finntip wide-orifice pipette tips. 10µL of the re-suspended nuclei with 990µL of PBS was then used for counting nuclei yield. The remaining 100 µL re-suspended nuclei were snap-frozen in liquid nitrogen and stored at -70*^◦^*C before being used for the PRO-seq nuclear-run-on reactions.

#### PRO-seq Across Species

PRO-seq datasets were prepared as described in [60]. Between 3-16 million nuclei per dataset were used for PRO-seq transcription run-on. Additionally, the run-on reaction was ran with a mixture of rNTP and Biotin-11-CTP (Biotin-11-CTP at 0.025 mM from PerkinElmer Ref. NEL542001EA; rCTP at 0.025 mM from Promega Ref. E604B, rATP at 0.125 mM Ref. E601B, rGTP at 0.125 mM Ref. E603B, and rUTP at 0.125 mM Ref. E6021). 1% of S2 Drosophila melanogaster nuclei relative to the number of the sample nuclei were added during the run-on reaction as a normalization spike-in. Total RNA was extracted using a phenol/chloroform precipitation. Isolated RNA was fragmented using base hydrolysis with NaOH. Biotinylated fragmented nascent transcripts were isolated using a first streptavidin Dynabeads M-280 (Invitrogen Ref. 11206D) pull down, and the VRA3 RNA adaptor was ligated at their 3*^′^* end. A second streptavidin bead pull-down was performed, followed by the enzymatic modifications of the RNA fragment 5*^′^* ends with a pyrophosphohydrolase and a polynucleotide kinase, and the VRA5 RNA adaptor was ligated at their fixed 5*^′^*ends. A third streptavidin bead pull-down was performed, followed by the reverse transcription of the resulting adaptor-ligated libraries. The libraries were cleaned up with AMPure XP beads (Beckman Coulter Ref. A63881). Then, the libraries were amplified using 13 PCR cycles and cleaned up again with another round of AMPure XP beads. The resulting library concentrations were measured with the Qubit dsDNA high sensitivity assay (Invitrogen Ref. Q32851), and their size distributions assessed using the Agilent High Sensitivity D1000 ScreenTape.

#### Data Processing Across Species

Cross species PRO-seq datasets were processed using the Nextflow pipeline found in https://github.com/Dowell-Lab/Nascent-Flow. Read quality was assessed using FastQC v0.11.5 for human, gibbon, rhesus, and squirrel monkey samples and version 0.11.8 for mouse samples. The human genome (hg38), gibbon reference genome (nomLeu3), rhesus genome (rheMac10), squirrel monkey genome (saiBol1), and mouse genome (mm10) were obtained from UCSC goldenPath. For squirrel monkey, contigs numbered from JH378105 to JH378420, were renamed as chr1 to chr316, respectively.

#### Arsenic Experiment

Human osteosarcoma U-2 OS (U-2 OS cells provided by Paul Anderson’s Lab at Brigham and Women’s Hospital, Boston, MA, USA [61]) cells were maintained in DMEM with 10% FBS and 1% penicillin/streptomycin at 37*^◦^*C, 5% CO_2_. For each PRO-seq condition (untreated and stressed), 20 million cells were seeded in five 15 cm plates. After 24 hours, to induce stress, cells were treated with NaAsO_2_ (S7400; Sigma-Aldrich; 500 *µ*M) for 1 hour prior to nuclei isolation. Nuclei isolation prep was done as mentioned in the section above (Cell Conditions and Nuclei Isolation Across Species), except that the nuclei counting step was omitted, and all nuclei were used.

#### PRO-seq Arsenic Experiment

PRO-seq datasets in the arsenic experiment were prepared as described in [60] (see PRO-seq Across Species above for more details), with three deviations. First, the runon reaction spanned 10 minutes instead of 3 minutes. Second, 3.5 million S2 Drosophila melanogaster nuclei were added before the run-on reaction as a normalization spike-in. Third, the libraries were amplified using 15 PCR cycles.

#### Data Processing Arsenic Experiment

PRO-seq datasets from the arsenic experiment were also processed using the Nextflow pipeline found in https://github.com/Dowell-Lab/Nascent-Flow. Read quality was assessed using FastQC (v0.11.8).

#### Data Availability

| Description | Accession Number(s) |
| --- | --- |
| Arsenic PRO-seq control experiments | GSM9321236, GSM9321237 |
| Arsenic PRO-seq perturbed experiments | GSM9321238, GSM9321239 |
| Human PRO-seq experiments | GSM6716757, GSM6716758 |
| Gibbon monkey PRO-seq experiments | GSM6716755, GSM6716756 |
| Rhesus monkey PRO-seq experiments | GSM6716759, GSM6716760 |
| Squirrel monkey PRO-seq experiments | GSM6716761, GSM6716762 |
| ATAC-seq experiment | GSM5269423 |

#### Curation of Isolated Gene Lists

The LIET model parametrizes transcription at a single gene. To avoid overlapping transcription and to provide adequate depth per gene, we created a computational pipeline to identify transcriptionally isolated genes on a per-cell-type basis. Details of this pipeline are at https://github.com/Dowell-Lab/Disassociation Analysis/tree/ main/Gene-Curation. The computational pipeline for identifying transcriptionally isolated genes began with an annotation filter, where genes with MANE (version 1.4) [62] annotations within 3kb upstream or 20kb downstream were removed, along with their overlapping neighbors. Additionally, we added two manually curated gene sets to our initial list prior to filtering: the set previously published in Stanley et al. 2025 [26] and a new set curated using the same criteria as Stanley et al. 2025 [26]. By adding these lists, we expanded the number of genes assessed. Coverage (0.001x per gene) and complexity (unique reads/total reads) filters (0.001x) were applied, followed by a gene length filter, as we found LIET performs poorly on short genes *<*7kb long. Coverage was calculated on MANE transcript annotations (version 1.4), where the 5*′* end of the transcript annotation was truncated (1/3rd of the gene length subtracted from the 5*′* end, with a max value of 750bps) to prevent all of the coverage being present at the 5*′* end in the loading and initiation step.

After fitting the filtered genes with LIET, genes with high error residuals and no termination component (0% of the reads to termination or *>*95% of the reads to a single phase of transcription) were considered poor fits and removed. Any genes with antisense transcription (0.005x or more on the antisense strand *±*3kb around *µ_T_*) or with a bidirectional *±*5kb around *µ_T_* (identified by Tfit [63]) were also removed to avoid confounding *µ_T_* values with unannotated bidirectionals. Finally, genes were then filtered to contain a coverage of *>* 0.001x *±*3kb around *µ_T_*, to ensure sufficient depth at the disassociation peak for model fitting. We increased the coverage cutoff to be *>* 0.0075x *±*3kb around *µ_T_* to account for low data depth and complexity in the arsenic control and experimental samples, as manual inspection indicated lower complexity within these samples.

We identified shared orthologs between human and each target species (rhesus, gibbon, squirrel monkey, and mouse) using Benchmarking Universal Single-Copy Orthologs (BUSCO) version 5.8.1 [34]. Then, we intersected human gene annotations (RefSeq GCF_000001405.40-RS_2023_03) with the BUSCO output, selecting orthologs with *>*75% overlap in genomic coordinates with annotated genes. Finally, we lifted over MANE-selected human coordinates to each target species’ genome using the Progressive Cactus multiple-genome aligner [35, 36], allowing us to obtain annotations for common orthologs in the rhesus monkey, gibbon monkey, squirrel monkey, and mouse genomes. After performing the lift over, fragments within 5000bp of each other were merged to consolidate fragmented regions (per-gene). Only the longest merged region was kept for each gene.

We applied the same gene curation steps used for human cell type data (meta-sample generation and coverage/bidirectional filtering) to each species. However, unlike the human analysis, we omitted the annotation-isolation filter to avoid excessively restricting the number of genes we analyzed. Additionally, the mouse sample was filtered at a higher coverage cutoff for the gene body (0.02x) prior to running LIET. After gene curation, we ran LIET on each species’ gene set independently. Specifics on ortholog selection and how we used the Progressive Cactus multiple-genome aligner are on our GitHub: https://github.com/Dowell-Lab/Disassociation Analysis/tree/main/ Cross-Species-liftOver.

#### LIET Model Inputs and Prior Selection

The LIET software (v1.0.0) allows the user to define 5*^′^* and 3*^′^* end reference points for each gene (*z*_5_,*z*_3_). These positions are used primarily to reduce runtime, as unbounded searches for parameterization with LIET is computationally expensive. MANE annotations for the transcription start site and the 5*^′^* end of the UTR were used for *z*_5_ and *z*_3_, respectively [62]. *z*_3_ is selected in this manner to allow for more genomic coordinate space downstream of elongation to identify a disassociation peak. Notably, *µ_T_* is consistently called in roughly the same genomic coordinate space regardless of what value *z*_3_ is given, as long as *z*_3_ is near the 3*^′^* end of the gene and does not proceed past the 3*^′^*end slowing peak (Supplemental Figure S19A-B). This paper employed the same priors recommended in the LIET model paper [26].

It is also worth noting that the 3*^′^* end of each read was used for the LIET input for maximum accuracy on the position of disassociation peak [26]. The Nextflow pipeline used to create these 3*′* bedGraphs is located on our GitHub repository:https://github.com/Dowell-Lab/Disassociation_Analysis/tree/main/3p Bedgraph_Generation.

Additionally, we used 3kb 5*′* pad values for the LIET input for all genes assessed. The 3*′* pad was 20kb for samples in control conditions and 30kb for samples prepared under conditions that induce run-on, providing extra genomic coordinate space to capture run-on transcription.

#### Data Analysis

A Jupyter Notebook containing the data analysis for this manuscript and a file with all software versions can be found on our GitHub: https://github.com/Dowell-Lab/ Disassociation Analysis/tree/main/Data-Analysis.

Briefly, we quantified technical variation (i.e., measurement variation) between human cell types and replicates (Figures 1C and D) by fitting the reproducibility of *|µ_T_ −* TCS| values (between samples) to a Normal distribution. We then set the measurement error as the 95% confidence interval, labeled in *|µ_T_ −* TCS| scatter plots as the shaded grey bar off the 1:1 line. When comparing across cell types, we create a single meta-sample per cell type by combining all replicates available. For cross-species comparisons (i.e., human vs. rhesus monkey, Figure 3D), we measured technical variation by calculating within-species measurement variation on *|µ_T_ − TCS|* across each replicate comparison (i.e., 2*σ* = 2993bp for human, 2*σ* = 3159bp for rhesus). We then utilize the larger of the two bounds when assessing the cross-species comparison (2*σ* = 3159). For the cross-perturbation analyses, we measured measurement variation by first plotting *|µ_T_ − TCS|* between control replicates for a given experiment. We then calculated 2*σ* from the 1:1 line between the replicates for each species and applied these bounds to the control vs. perturbation comparison.

We utilize the 95% confidence interval (2*σ*) as a classifier for identifying perturbation-induced shifts in disassociation on a per-gene basis. In other words, only differences larger than 2*σ* are considered changes in disassociation. Using the set of genes with changes to disassociation, we then utilize a binomial test to assess whether the number of genes outside the confidence interval exceeds expectation (5% of values would fall outside the 95% confidence interval). We used the binom.sf function (part of the scipy.stats.binom package) for these binomial tests. We used a 1-sided test to look for alterations that shift disassociation downstream (run-on) or upstream (shortening). Moreover, we used a 2-sided test to detect alterations that shift disassociation in either direction (outside the 2*σ* bounds).

We test for enrichment of the upstream T-rich and GC-rich subsets in the set of genes with altered disassociation (run-on or shortening) by using a hypergeometric test with the hypergeom.sf function (part of the scipy.stats.hypergeom package). Hypergeometric test results on genes below 2*σ* under HSV-1 infection, not referenced in the main figures, are shown in Supplemental Figure S18C.

Base composition plots in Figures 2-3 were generated by averaging the A, T, G, and C content at every position for a fixed window (*µ_T_ ±*4*kb*) across all genes investigated. We evenly distributed unknown bases (N’s) across all four bases before averaging. We normalized these counts by the total number of sequences to calculate the proportion of each base at each position. Base composition plots at *µ_T_* across perturbations that are not referenced in the Results section are shown in Supplemental Figure S13D-F.

Sequence alignments in Figure 3 were generated using AlignIO from the Biopython package. Each position was assigned a specific color indicating whether it represented a nucleotide insertion, deletion, or match. We then plotted the alignment between the two species (rhesus monkey and human) utilizing this color scheme.

Metagene plots in Figure 4 were generated by calculating PRO-seq coverage 1kb upstream and 7kb downstream of the TCS. We used a sliding-window approach, as previously used on our CDK7 inhibition work [44]. We used MANE annotated transcript coordinates to obtain the TCS [62]. Minus-strand genes were reverse-complemented before averaging. We only included regions with a total coverage exceeding 50 reads per bin. Additionally, reads were counted across both strands. The metagenes represent the mean of binned coverage across all regions in control and perturbed conditions.

## Author Contribution

G.E.F.B. ran LIET, analyzed all data, generated all figures, and co-wrote the manuscript. J.T.S. developed the LIET model and contributed to the study design. D.R. generated, mapped, and quality-controlled PRO-seq data across primates. J.F.C. generated, mapped, and quality-controlled ATAC-seq data. N.R. and R.P. generated, mapped, and quality-control-checked PRO-seq data exposed to arsenic. M.A.A. organized the study and interpreted the results. R.D.D. organized the study, interpreted results, and co-wrote the manuscript.

## Acknowledgments

We thank Jen Kugel for consultation on run-on transcription. We also thank Dylan Taatjes for feedback on the manuscript draft and consultation on kinase activity. Additionally, we express our gratitude to Alejandro Juarros for the manual inspection of LIET fits under run-on-inducing conditions. Moreover, we thank Andrew Sanders for his help with gene curation. We are also grateful to the BioFrontiers IT department. Their support in maintaining the HPC system enabled our work on this project.

## References

[1] Proudfoot, N.J.: Transcriptional termination in mammals: Stopping the rna polymerase ii juggernaut. Science 352(6291), 9926 (2016)

[2] West, S., Proudfoot, N.J.: Transcriptional termination enhances protein expression in human cells. Molecular Cell 33(3), 354–364 (2009)

[3] Au, V., Li-Leger, E., Raymant, G., Flibotte, S., Chen, G., Martin, K., Fernando, L., Doell, C., Rosell, F.I., Wang, S., et al.: CRISPR/Cas9 methodology for the generation of knockout deletions in Caenorhabditis elegans. G3: Genes, Genomes, Genetics 9(1), 135–144 (2019)

[4] Ryder, E., Gleeson, D., Sethi, D., Vyas, S., Miklejewska, E., Dalvi, P., Habib, B., Cook, R., Hardy, M., Jhaveri, K., et al.: Molecular characterization of mutant mouse strains generated from the EUCOMM/KOMP-CSD ES cell resource. Mammalian Genome 24, 286–294 (2013)

[5] Rosa-Mercado, N.A., Steitz, J.A.: Who let the dogs out? biogenesis of stress-induced readthrough transcripts. Trends in Biochemical Sciences 47(3), 206–217 (2022)

[6] Morgan, M., Shiekhattar, R., Shilatifard, A., Lauberth, S.M.: It’s a DoG-eat-DoG world—altered transcriptional mechanisms drive downstream-of-gene (DoG) transcript production. Molecular cell (2022)

[7] Vilborg, A., Steitz, J.A.: Readthrough transcription: How are DoGs made and what do they do? RNA biology 14(5), 632–636 (2017)

[8] Vilborg, A., Passarelli, M.C., Yario, T.A., Tycowski, K.T., Steitz, J.A.: Widespread inducible transcription downstream of human genes. Molecular Cell 59(3), 449–461 (2015)

[9] Wiesel, Y., Sabath, N., Shalgi, R.: DoGFinder: a software for the discovery and quantification of readthrough transcripts from RNA-seq. BMC Genomics 19(1), 597 (2018)

[10] Rutkowski, A.J., Erhard, F., L’Hernault, A., Bonfert, T., Schilhabel, M., Crump, C., Rosenstiel, P., Efstathiou, S., Zimmer, R., Friedel, C.C., et al.: Widespread disruption of host transcription termination in HSV-1 infection. Nature communications 6(1), 7126 (2015)

[11] Bauer, D.L., Tellier, M., Martínez-Alonso, M., Nojima, T., Proudfoot, N.J., Murphy, S., Fodor, E.: Influenza virus mounts a two-pronged attack on host RNA polymerase II transcription. Cell Reports 23(7), 2119–2129 (2018)

[12] Zhang, H., Rigo, F., Martinson, H.G.: Signal-dependent transcription termination occurs through a conformational change mechanism that does not require cleavage at the poly (a) site. Mol Cell 59(3), 437–48 (2015)

[13] Han, Z., Moore, G.A., Mitter, R., Martinez, D.L., Wan, L., Svejstrup, A.B.D., Rueda, D.S., Svejstrup, J.Q.: Dna-directed termination of rna polymerase ii transcription. Molecular cell 83(18), 3253–3267 (2023)

[14] Davidson, L., Rouvière, J.O., Sousa-Luís, R., Nojima, T., Proudfoot, N.J., Jensen, T.H., West, S.: Dna-directed termination of mammalian rna polymerase ii. Genes & Development 38(21-24), 998–1019 (2024)

15. [15] Kuehner, J.N., Pearson, E.L., Moore, C.: Unravelling the means to an end: RNA polymerase II transcription termination. Nature reviews Molecular cell biology 12(5), 283–294 (2011)

[16] Eaton, J.D., Francis, L., Davidson, L., West, S.: A unified allosteric/torpedo mechanism for transcriptional termination on human protein-coding genes. Genes Dev 34(1-2), 132–145 (2020)

[17] Eaton, J.D., West, S.: Termination of transcription by RNA polymerase II: BOOM! Trends in Genetics 36(9), 664–675 (2020)

[18] Fong, N., Brannan, K., Erickson, B., Kim, H., Cortazar, M.A., Sheridan, R.M., Nguyen, T., Karp, S., Bentley, D.L.: Effects of transcription elongation rate and xrn2 exonuclease activity on rna polymerase ii termination suggest widespread kinetic competition. Molecular cell 60(2), 256–267 (2015)

[19] West, S., Gromak, N., Proudfoot, N.J.: Human 5’ -¿ 3’ exonuclease xrn2 promotes transcription termination at co-transcriptional cleavage sites. Nature 432(7016), 522–525 (2004)

[20] Mahat, D.B., Kwak, H., Booth, G.T., Jonkers, I.H., Danko, C.G., Patel, R.K., Waters, C.T., Munson, K., Core, L.J., Lis, J.T.: Base-pair-resolution genome-wide mapping of active rna polymerases using precision nuclear run-on (pro-seq). Nat. Protoc. 11(8), 1455–1476 (2016)

[21] Wissink, E.M., Vihervaara, A., Tippens, N.D., Lis, J.T.: Nascent RNA analyses: tracking transcription and its regulation. Nature Reviews Genetics 20(12), 705– 723 (2019)

[22] Azofeifa, J., Allen, M.A., Lladser, M., Dowell, R.: FStitch: A fast and simple algorithm for detecting nascent RNA transcripts. Proceedings of the 5th ACM Conference on Bioinformatics, Computational Biology, and Health Informatics, 174–183 (2014) 10.1145/2649387.2649427

[23] Chae, M., Danko, C.G., Kraus, W.L.: groHMM: a computational tool for identifying unannotated and cell type-specific transcription units from global run-on sequencing data. BMC Bioinformatics 16(1), 222 (2015)

[24] Schwalb, B., Michel, M., Zacher, B., Frühauf, K., Demel, C., Tresch, A., Gagneur, J., Cramer, P.: Tt-seq maps the human transient transcriptome. Science 352(6290), 1225–1228 (2016)

[25] Zhao, Y., Liu, L., Hassett, R., Siepel, A.: Model-based characterization of the equilibrium dynamics of transcription initiation and promoter-proximal pausing in human cells. Nucleic Acids Res 51(21), 106 (2023)

[26] Stanley, J.T., Barone, G.E.F., Townsend, H.A., Sigauke, R.F., Allen, M.A., Dowell, R.D.: LIET model: capturing the kinetics of RNA polymerase from loading to termination. Nucleic Acids Res 53(7), 246 (2025)

[27] Mayer, A., Landry, H.M., Churchman, L.S.: Pause & go: from the discovery of RNA polymerase pausing to its functional implications. Curr Opin Cell Biol 46, 72–80 (2017)

[28] Sigauke, R.F., Sanford, L., Maas, Z.L., Jones, T., Stanley, J.T., Townsend, H.A., Allen, M.A., Dowell, R.D.: Atlas of nascent RNA transcripts reveals tissue-specific enhancer to gene linkages. BMC Genomics 26(1), 406 (2025)

[29] Kopczyńska, M., Saha, U., Romanenko, A., Nojima, T., Gdula, M.R., Kamieniarz-Gdula, K.: Defining gene ends: Rna polymerase ii ctd threonine 4 phosphorylation marks transcription termination regions genome-wide. Nucleic Acids Research 53(2), 1240 (2025) 10.1093/nar/gkae1240

[30] Hintermair, C., Heidemann, M., Koch, F., Descostes, N., Gut, M., Gut, I., Fenouil, R., Ferrier, P., Flatley, A., Kremmer, E., Chapman, R.D., Andrau, J.-C., Eick, D.: Threonine-4 of mammalian rna polymerase ii ctd is targeted by polo-like kinase 3 and required for transcriptional elongation. The EMBO journal 31(12), 2784–2797 (2012)

[31] Etchegaray, J.-P., Zhong, L., Li, C., Henriques, T., Ablondi, E., Nakadai, T., Rechem, C.V., et al.: The histone deacetylase sirt6 restrains transcription elongation via promoter-proximal pausing. Molecular Cell 75(4), 683–699 (2019)

[32] Breschi, A., Gingeras, T.R., Guigó, R.: Comparative transcriptomics in human and mouse. Nature Reviews Genetics 18(7), 425–440 (2017)

[33] Yue, F., Cheng, Y., Breschi, A., Vierstra, J., Wu, W., Ryba, T., Sandstrom, R., et al.: A comparative encyclopedia of dna elements in the mouse genome. Nature 515(7527), 355–364 (2014)

[34] Manni, M., Berkeley, M.R., Seppey, M., Simão, F.A., Zdobnov, E.M.: Busco update: Novel and streamlined workflows along with broader and deeper phylogenetic coverage for scoring of eukaryotic, prokaryotic, and viral genomes. Molecular Biology and Evolution 38(10), 4647–4654 (2021) 10.1093/molbev/msab199

[35] Zoonomia Consortium: A comparative genomics multitool for scientific discovery and conservation. Nature 587, 240–245 (2020) 10.1038/ s41586-020-2876-6

[36] Kuderna, L.F.K., Ulirsch, J.C., Rashid, S., et al.: Identification of constrained sequence elements across 239 primate genomes. Nature 625, 735–742 (2024) 10.1038/s41586-023-06798-8

[37] Cardiello, J.F., Westfall, J., Dowell, R., Allen, M.A.: Characterizing primary transcriptional responses to short term heat shock in down syndrome. PLOS ONE 19(8), 1–18 (2024) 10.1371/journal.pone.0307375

[38] Hennig, T., Michalski, M., Rutkowski, A.J., Djakovic, L., Whisnant, A.W., Friedl, M.-S., Jha, B.A., Baptista, M.A.P., L’Hernault, A., Erhard, F., Dölken, L., Friedel, C.C.: HSV-1-induced disruption of transcription termination resembles a cellular stress response but selectively increases chromatin accessibility downstream of genes. PLOS Pathogens 14(3), 1–27 (2018)

[39] Wang, X., Hennig, T., Whisnant, A., Erhard, F., Prusty, B., Friedel, C., Forouzmand, E., et al.: Herpes simplex virus blocks host transcription termination via the bimodal activities of icp27. Nat. Commun. 11, 293 (2020)

[40] Birkenheuer, C.H., Danko, C.G., Baines, J.D.: Herpes simplex virus 1 dramatically alters loading and positioning of RNA polymerase II on host genes early in infection. Journal of Virology 92(8), 02184–17 (2018) 10.1128/JVI.02184-17. Accessed 2018-05-23

[41] Steinparzer, I., Sedlyarov, V., Rubin, J.D., Eislmayr, K., Galbraith, M.D., Levandowski, C.B., Vcelkova, T., Sneezum, L., Wascher, F., Amman, F., et al.: Transcriptional responses to IFN-*γ* require mediator kinase-dependent pause release and mechanistically distinct CDK8 and CDK19 functions. Molecular cell 76(3), 485–499 (2019)

[42] Birkenheuer, C.H., Danko, C.G., Baines, J.D.: Herpes simplex virus 1 dramatically alters loading and positioning of rna polymerase ii on host genes early in infection. Journal of Virology 92(10), 02184–17 (2018) 10.1128/JVI.02184-17

[43] Jones, T., Feng, J., Luyties, O., Cozzolino, K., Sanford, L., Rimel, J.K., Ebmeier, C.C., Shelby, G.S., Watts, L.P., Rodino, J., Rajagopal, N., Hu, S., Brennan, F., Maas, Z.L., Alnemy, S., Richter, W.F., Koh, A.F., Cronin, N.B., Madduri, A., Das, J., Cooper, E., Hamman, K.B., Chuaqui, C., Carulli, J.P., Allen, M.A., Spencer, S., Kotecha, A., Marineau, J.J., Greber, B.J., Dowell, R.D., Taatjes, D.J.: TFIIH kinase CDK7 drives cell proliferation through a common core transcription factor network. Science Advances 11(9), 9660 (2025) 10.1126/sciadv.adr9660

[44] Luyties, O., Sanford, L., Rodino, J., Nagel, M., Jones, T., Rimel, J.K., Ebmeier, C.C., Palacio, M., Shelby, G.S., Cozzolino, K., Brennan, F.: Multi-omics and biochemical reconstitution reveal CDK7-dependent mechanisms controlling RNA polymerase II function at gene 5*^′^*-and 3*^′^* ends. Cell Reports 44(7) (2025)

[45] Jaeger, M.G., Schwalb, B., Mackowiak, S.D., Velychko, T., Hanzl, A., Imrichova, H., Brand, M., Agerer, B., Chorn, S., Nabet, B., Ferguson, F.M., Müller, A., Bergthaler, A., Gray, N.S., Bradner, J.E., Bock, C., Hnisz, D., Cramer, P., Winter, G.E.: Selective mediator dependence of cell-type-specifying transcription. Nature Genetics 52(7), 719–727 (2020)

[46] Luyties, O., Taatjes, D.J.: The mediator kinase module: an interface between cell signaling and transcription. Trends Biochem. Sci. 47(4), 314–327 (2022)

[47] Sun, R., Fisher, R.P.: The cdk9-spt5 axis in control of transcription elongation by rnapii. J. Mol. Biol. 437(1), 168746 (2025)

[48] Tellier, M., Zaborowska, J., Neve, J., Nojima, T., Hester, S., Fournier, M., Furger, A., Murphy, S.: Cdk9 and pp2a regulate rna polymerase ii transcription termination and coupled rna maturation. EMBO reports 23(10), 54520 (2022)

[49] Laitem, C., Zaborowska, J., Isa, N.F., Kufs, J., Dienstbier, M., Murphy, S.: CDK9 inhibitors define elongation checkpoints at both ends of RNA polymerase II– transcribed genes. Nature structural & molecular biology 22(5), 396–403 (2015)

[50] Sansó, M., Levin, R.S., Lipp, J.J., Wang, V.Y.-F., Greifenberg, A.K., Quezada, E.M., Ali, A., Ghosh, A., Larochelle, S., Rana, T.M., Geyer, M., Tong, L., Shokat, K.M., Fisher, R.P.: P-TEFb regulation of transcription termination factor Xrn2 revealed by a chemical genetic screen for Cdk9 substrates 30(1), 117–131 (2016)

[51] Gupta, A., Sasse, S.K., Gruca, M.A., Sanford, L., Dowell, R.D., Gerber, A.N.: Deconvolution of multiplexed transcriptional responses to wood smoke particles defines rapid aryl hydrocarbon receptor signaling dynamics. J Biol Chem 297(4), 101147 (2021)

[52] Jonkers, I., Kwak, H., Lis, J.T.: Genome-wide dynamics of pol II elongation and its interplay with promoter proximal pausing, chromatin, and exons. eLife 3 (2014)

[53] Cardiello, J.F., Goodrich, J.A., Kugel, J.F.: Heat shock causes a reversible increase in RNA polymerase II occupancy downstream of mRNA genes, consistent with a global loss in transcriptional termination. Molecular and Cellular Biology 38(18), 00181–18 (2018)

[54] Mellor, J., Woloszczuk, R., Howe, F.S.: The interleaved genome. Trends in Genetics 32(1), 57–71 (2016)

[55] Chen, C.-H., Pan, C.-Y., Lin, W.-c.: Overlapping protein-coding genes in human genome and their coincidental expression in tissues. Scientific Reports 9(1), 1–10 (2019)

[56] Shearwin, K., Callen, B., Egan, J.: Transcriptional interference: a crash course. Trends in Genetics 21, 339 (2005)

[57] Hadar, S., Meller, A., Saida, N., Shalgi, R.: Stress-induced transcriptional readthrough into neighboring genes is linked to intron retention. iScience 25(12), 105543 (2022) 10.1016/j.isci.2022.105543

[58] Corces, M., Trevino, A., Hamilton, E., et al.: An improved atac-seq protocol reduces background and enables interrogation of frozen tissues. Nat. Methods 14, 959–962 (2017)

[59] Core, L.J., Waterfall, J.J., Lis, J.T.: Nascent RNA sequencing reveals widespread pausing and divergent initiation at human promoters. Science 322(5909), 1845–1848 (2008)

[60] Fant, C.B., Levandowski, C.B., Gupta, K., Maas, Z.L., Moir, J., Rubin, J.D., Sawyer, A., Esbin, M.N., Rimel, J.K., Luyties, O., Marr, M.T., Berger, I., Dowell, R.D., Taatjes, D.J.: TFIID enables RNA polymerase II promoter-proximal pausing. Molecular Cell 78(4), 785–7938 (2020)

[61] Kedersha, N., Panas, M.D., Achorn, C.A., Lyons, S., Tisdale, S., Hickman, T., Thomas, M., Lieberman, J., McInerney, G.M., Ivanov, P., Anderson, P.: G3bp-caprin1-usp10 complexes mediate stress granule condensation and associate with 40s subunits. Journal of Cell Biology 212(7), 845–860 (2016)

[62] Morales, J., Pujar, S., Loveland, J.E., Astashyn, A., Bennett, R., Berry, A., Cox, E., Davidson, C., Ermolaeva, O., Farrell, C.M., Fatima, R.: A joint ncbi and emblebi transcript set for clinical genomics and research. Nature 604(7905), 310–315 (2022) 10.1038/s41586-022-04558-8

[63] Azofeifa, J.G., Allen, M.A., Hendrix, J.R., Read, T., Rubin, J.D., Dowell, R.D.: Enhancer RNA profiling predicts transcription factor activity. Genome Research (2018) 10.1101/gr.225755.117

[64] Jones, T., Sigauke, R.F., Sanford, L., Taatjes, D.J., Allen, M.A., Dowell, R.D.: A transcription factor (tf) inference method that broadly measures tf activity and identifies mechanistically distinct tf networks. bioRxiv (2024) 10.1101/2024.03.15.585303

