## Supplemental Figures and Table for "Base Composition Influences the Position and Precision of RNA Polymerase II Disassociation in Basal and Perturbed Conditions"

December 1, 2025

[1] BioFrontiers Institute, University of Colorado, 3415 Colorado Ave., UCB 596, Boulder CO 80309 USA
[2] Department of Molecular, Cellular and Developmental Biology, University of Colorado, 1945 Colorado Ave, UCB 347, Boulder CO 80309 USA
[3] Department of Computer Science, University of Colorado, 1111 Engineering Drive, UCB 430, Boulder CO 80309 USA
[4] Department of Biochemistry, University of Colorado, 3415 Colorado Ave., 596 UCB, Boulder CO 80309 USA
[5] Howard Hughes Medical Institute, 4000 Jones Bridge Road, Chevy Chase MD 20815 USA

### Supplementary Figures

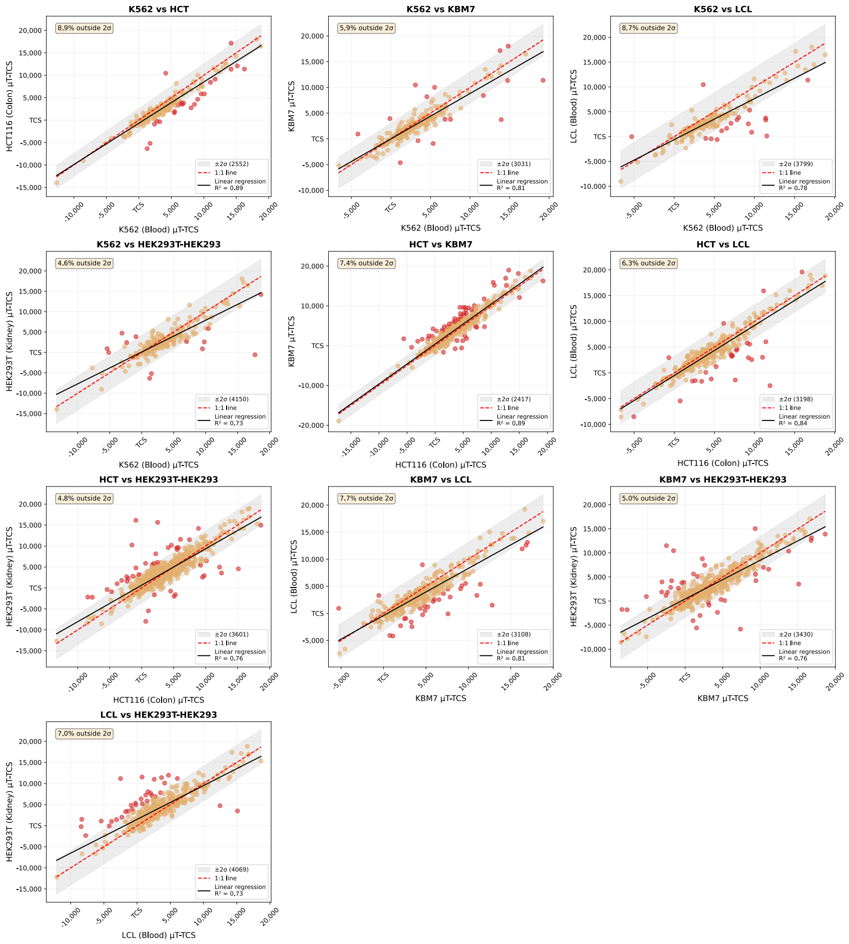


Figure S1: **Comparison of $|\mu_T - TCS|$ across human cell types.** 1:1 line (red dashed line) shows expected $\mu_T$ value if disassociation is perfectly reproducible between cell types. Measurement variation is captured by a Normal distribution, comparing human cell types (meta-samples, all replicates combined), with $2\sigma$ bounds (95% confidence interval) shown in grey on the scatter plots. The 95% confidence interval is calculated based on the variability in $|\mu_T - TCS|$ between the two cell types (on the meta-samples) for each subplot. Orange dots are genes contained within $2\sigma$ bounds, red dots are genes contained outside of $2\sigma$ bounds.

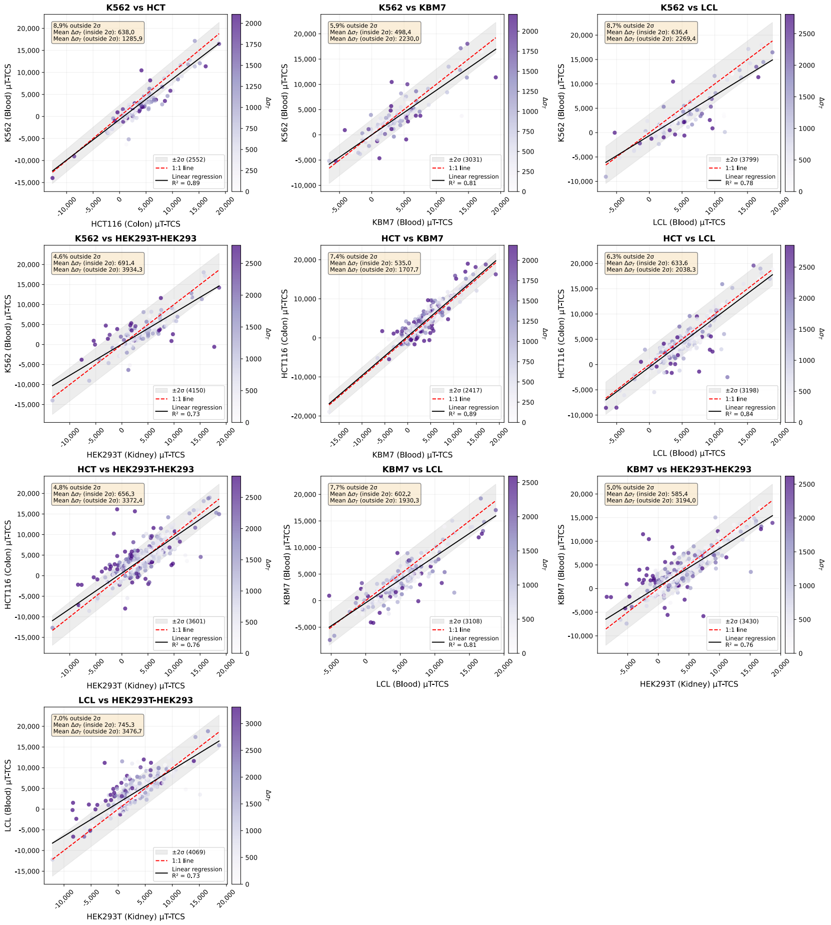


Figure S2: **Comparison of $|\mu_T - TCS|$ across human cell types, genes colored on changes in $\sigma_T$.** 1:1 line (red dashed line) shows expected $\mu_T$ value if disassociation is perfectly reproducible between cell types. We find the outlier genes (those outside the $2\sigma$ measurement variation gray zone) have higher $\sigma_T$ values (in purple, determined by LIET). Thus, as $\sigma_T$ becomes wider, $\mu_T$ becomes more variable– thus $\sigma_T$ accurately captures fidelity of the $\mu_T$ position.

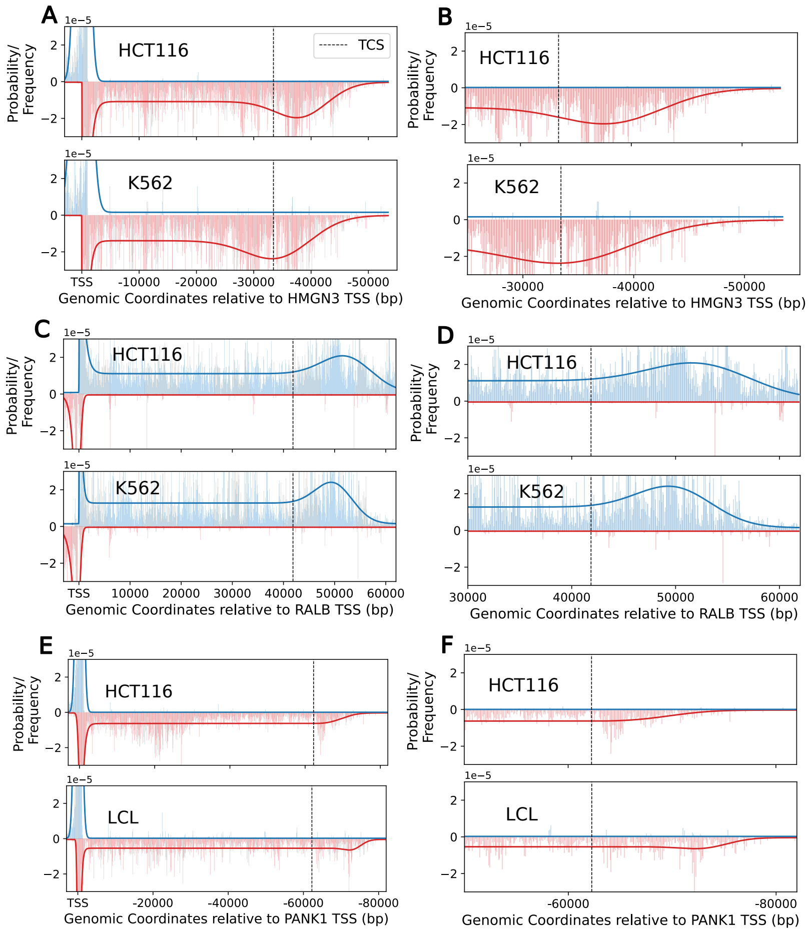


Figure S3: **LIET model fits for HMGN3 (A,B), RALB (C,D), and PANK1 (E,F) in different human cell types.** These genes exhibit $\mu_T$ values that deviate from expected technical variation. Wider $\sigma_T$ correlates to increased width in the disassociation peak. Data shown as histograms in strand color. Model fits as lines (blue: positive strand, red: negative strand). Dashed black vertical line marks TCS position.

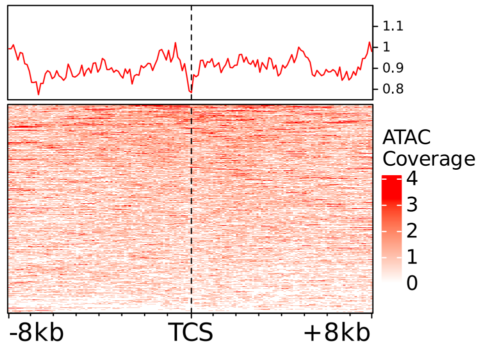


Figure S4: **Heatmap of ATAC-seq coverage ±8kb around the transcription cleavage site (TCS).** Rows reflect coverage at individual genes, top (red line) summarizes the heatmap as a meta-gene. Chromatin accessibility levels are higher at at $\mu_T$ relative to the TCS (Figure 1).

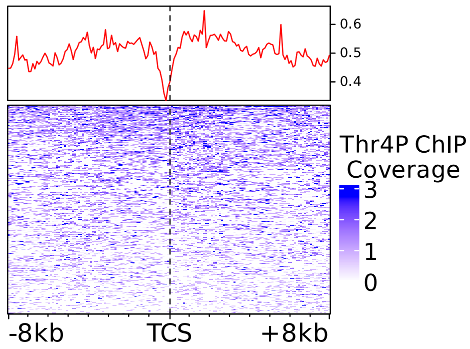


Figure S5: **Heatmap of Pol II CTD phosphorylation mark Th4P ChIP-seq coverage.** Window reflects ±8kb around the transcription cleavage site (TCS) [1]. Rows reflect coverage at individual genes, top (red line) summarizes the heatmap as a meta-gene. Th4P sharply decreases at this site and is enriched at $\mu_T$ (see Figure 1).

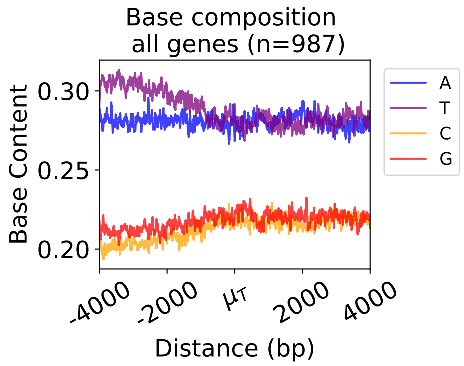


Figure S6: **Base composition ±4kb around $\mu_T$.** This plot is the averaged base composition at all genes (i.e., across both subplots in Figure 2A). Human LCL control sample prepped for primate comparisons (Supplementary Table 1). T content is enriched upstream of $\mu_T$ across all genes.

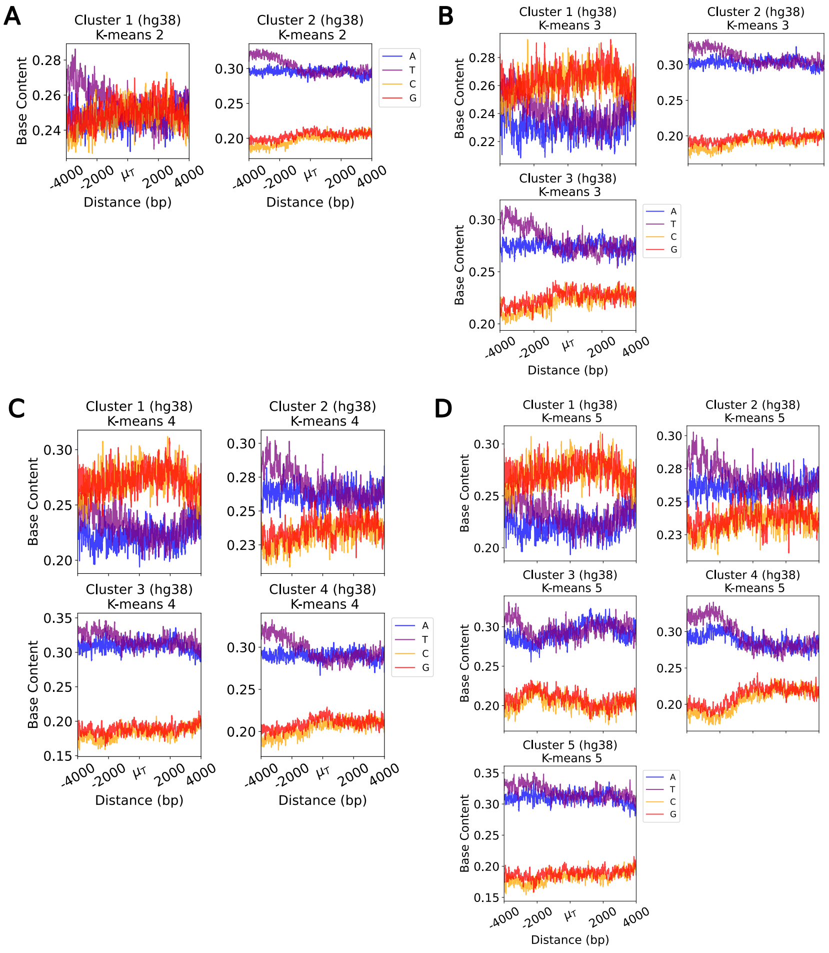


Figure S7: **Base composition patterns across k-means clusters in human LCL samples ±4kb around $\mu_T$.** K-means clustering was performed on AT content surrounding $\mu_T$ (±3kb). K-means of **A.** two, **B.** three, **C.** four, and **D.** five were computed. Generally two gene sets emerge: a GC-rich at $\mu\_T$, and a upstream T-rich set. Background base composition shown in Supplemental Figure S21.

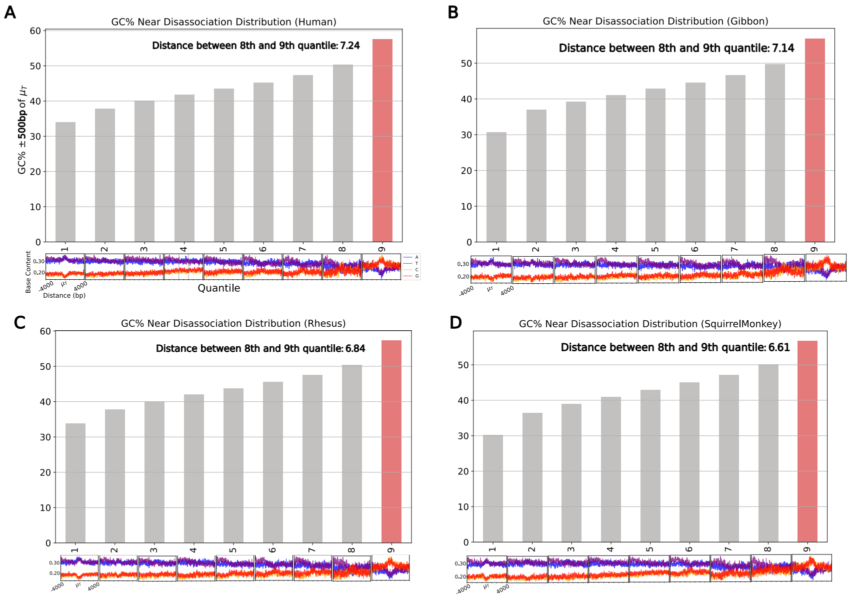


Figure S8: **Bargraphs of GC content ±1kb around $\mu_T$.** Shown are 9 quantiles, bars ordered by increasing GC content ±500$bp$ of $\mu_T$. Base composition plotted blow each quantile. Data shown for **A.** human, **B.** gibbon, **C.** rhesus, and **D.** squirrel monkey. In all cases, Q9 (in red) shows the strongest GC content and is used to define the GC-rich subset.

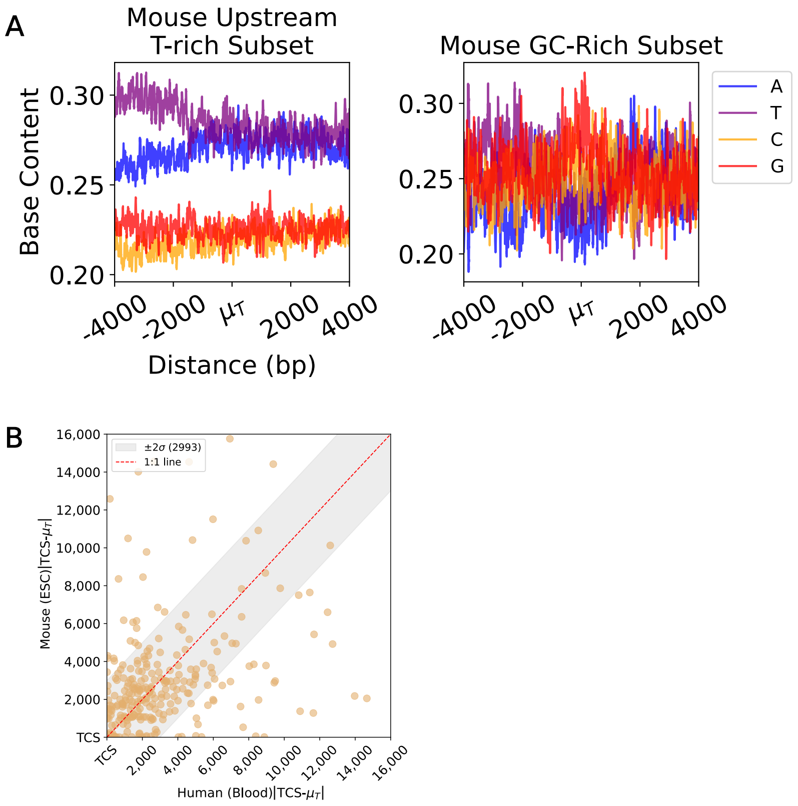


Figure S9: **Base composition around $\pm$4kb of $\mu_T$ (mouse ESCs) and $|\mu_T - TCS|$ (mouse vs. human). A.** Mouse (mm10) nascent sequencing data [2] shows the same base composition patterns $\pm$4kb around $\mu_T$ compared to human samples. **B.** $|\mu_T - TCS|$ in human cells (x-axis) vs. $|\mu_T - TCS|$ in mouse cells (y-axis) shows distance traveled after TCS varies between the species. $2\sigma$ bounds are defined by human replicates, as they are more variable relative to the mouse replicates.

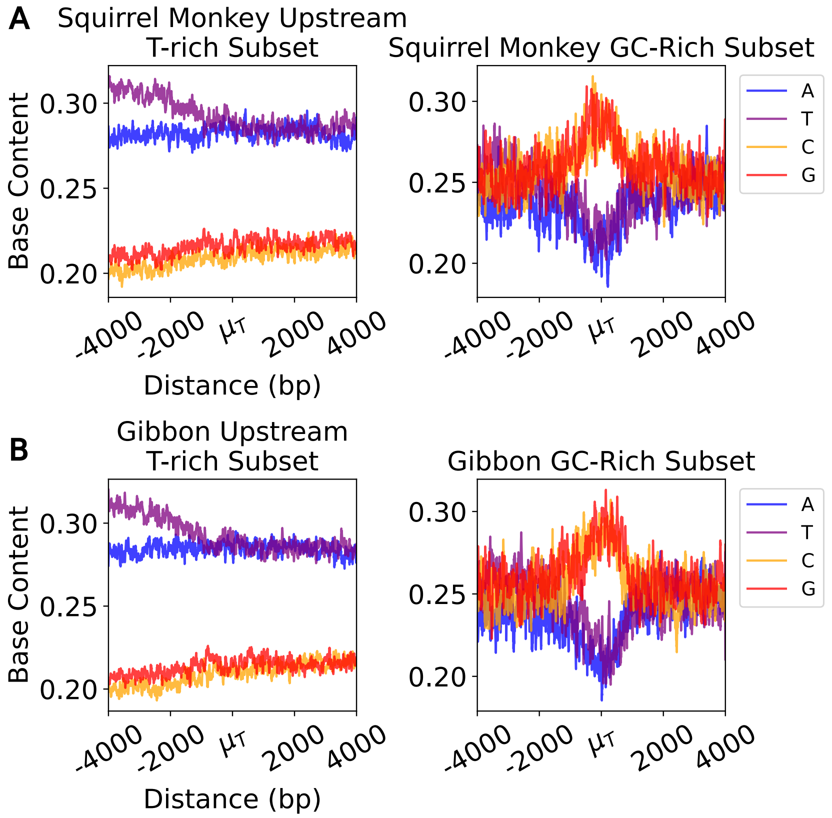


Figure S10: **GC-rich and upstream T-rich base composition patterns ±4kb around $\mu_T$ in rhesus and gibbon monkeys.** Base composition plots ±4kb around $\mu_T$ for **A.** rhesus and **B.** gibbon monkeys from quantile analysis (9 quantiles, ordered on GC content (±1kb around $\mu_T$). The upstream T-rich pattern is comprised of quantiles 1-8. GC-rich pattern is comprised of quantile 9 (see Supplemental Figure S8 for quantile plot).

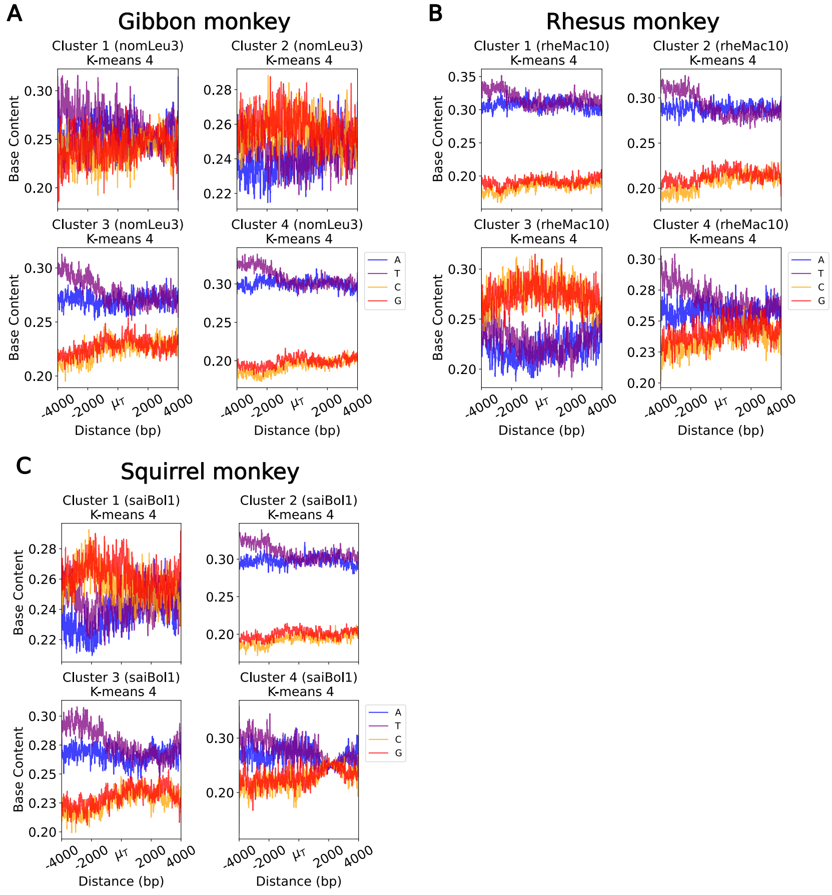


Figure S11: **Primate base composition patterns ±4kb around $\mu_T$ across k-means clusters.** K-means (k-means of 2-5) clustering was performed on AT content surrounding $\mu_T$ (±3kb). Only k-means of 4 is shown in **A.** gibbon, **B.** rhesus, and **C.** squirrel monkey for simplicity.

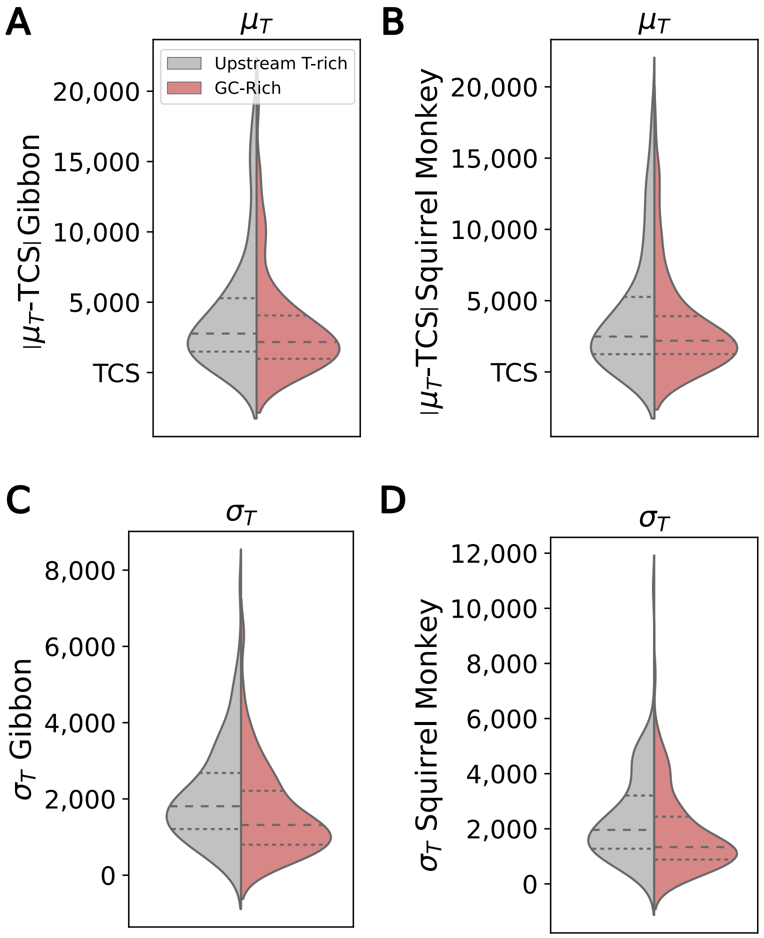


Figure S12: **Distributions of $|\mu_T - TCS|$ and $\sigma_T$ across species.** $|\mu_T - TCS|$ distribution in the upstream T-rich vs. GC-rich subsets in **A.** gibbon and **B.** squirrel monkeys. $\sigma_T$ distribution in the upstream T-rich vs. GC-rich subsets in **C.** gibbon monkey and **D.** squirrel monkey. The upstream T-rich subset was randomly sub-sampled to equal the same size as the GC-rich subset in each comparison.

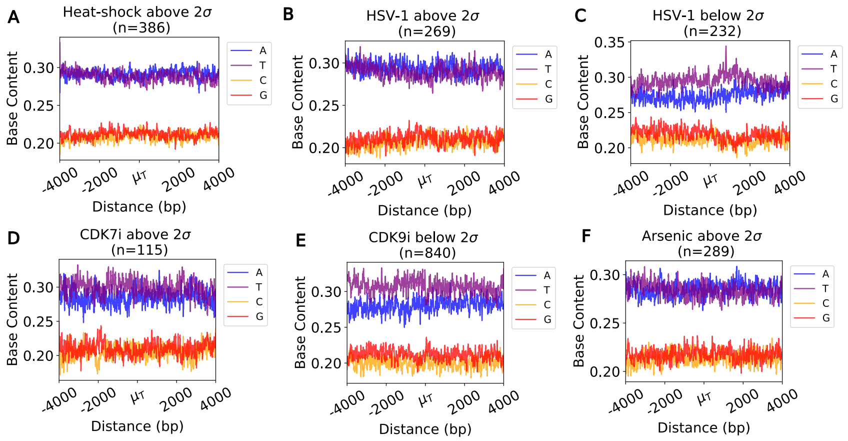


Figure S13: **Base composition pattern ±4kb around $\mu_T$ above or below 2$\sigma$ (depending on perturbation) upon stressor. A.** Heat-shock (above 2$\sigma$), **B.** HSV-1 (above 2$\sigma$), **C.** HSV-1 (below 2$\sigma$), **D.** CDK7 inhibition (above 2$\sigma$), **E.** CDK9 inhibition (below 2$\sigma$), and **F.** arsenic exposure (above 2$\sigma$)
.

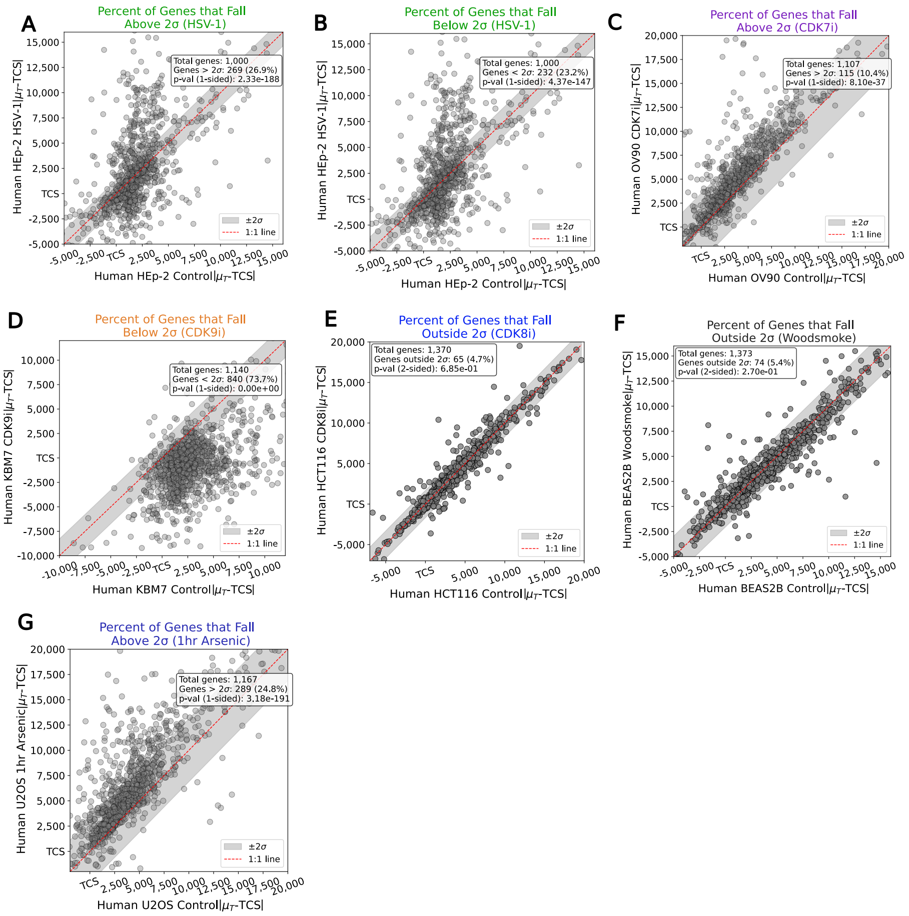


Figure S14: $|\mu_T - TCS|$ **in control vs perturbation.** Binomial test p-values assess enrichment of genes beyond $2\sigma$ bounds. One-sided tests examined genes above $2\sigma$ (run-on) or below $2\sigma$ ($3\prime$ end shortening); two-sided tests examined genes outside $2\sigma$ (general shift). **A.** HSV-1, run-on. **B.** HSV-1, $3\prime$ end shortening. **C.** CDK8, general shift. **D.** Wood smoke, general shift. **E.** CDK7 inhibition, run-on. **F.** CDK9 inhibition, $3\prime$ end shortening. **G.** Arsenic exposure, run-on.

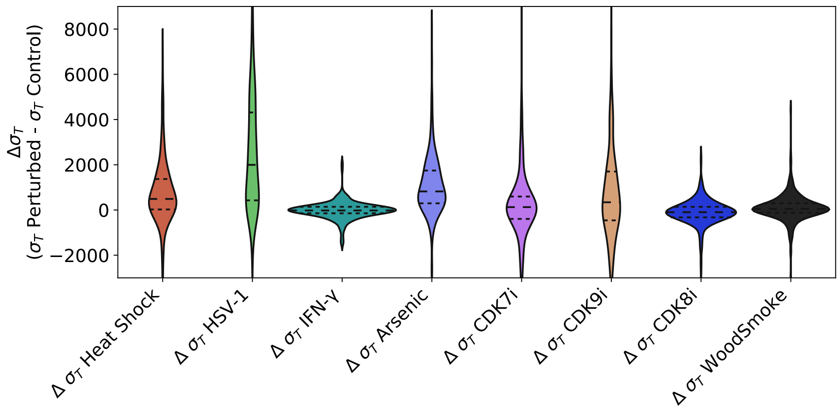


Figure S15: **Change ($\Delta$) in $\sigma_T$ across perturbations.** HSV-1 (green) has the strongest impact on Pol II disassociation precision. CDK9 inhibition (yellow) also negatively impacts disassociation precision. Heat-shock (orange), Arsenic (light blue), CDK7 inhibition (pink), CDK8 inhibition (dark blue) and wood smoke (black) have only limited impact on disassociation peak precision. IFN-$\gamma$ has the smallest impact on Pol II disassociation peak precision. $\Delta\sigma_T$ calculated as perturbed - control conditions.

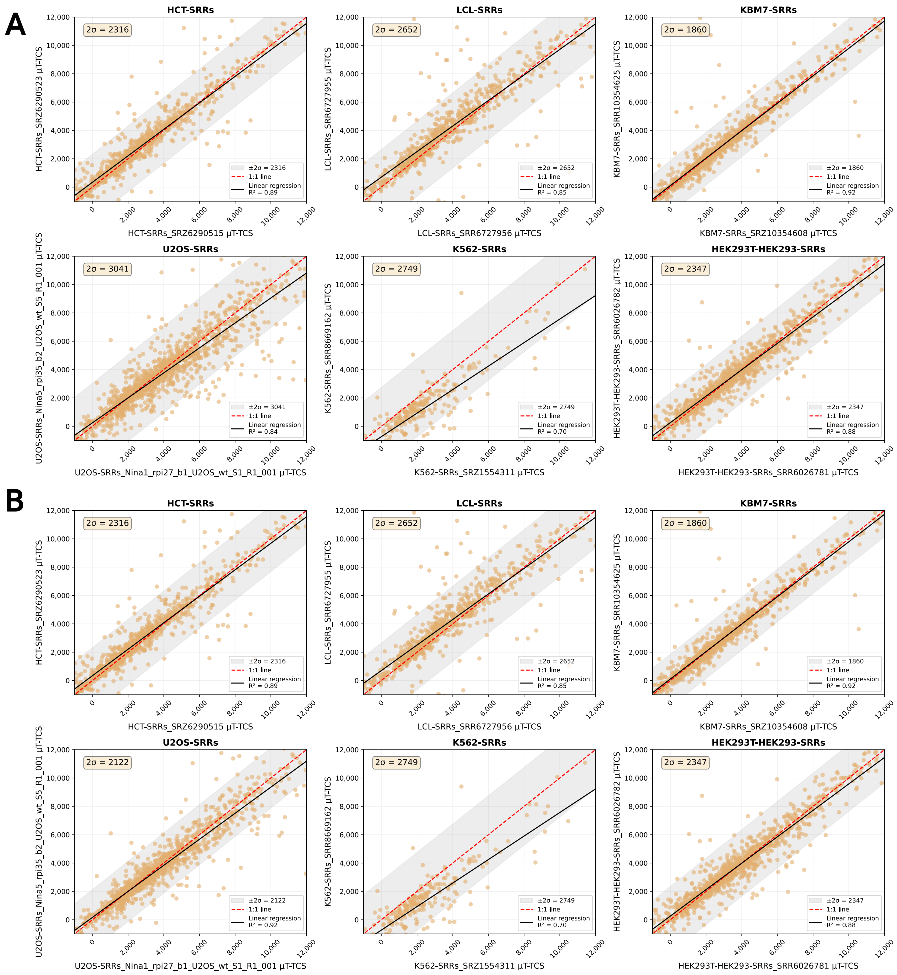

Figure S16: **Increasing the sense strand coverage filter (±3kb around $\mu_T$) in U-2 OS cells adjusts for low sequencing depth and complexity. A.** Comparison of $|\mu_T - TCS|$ between replicates across human cell types. U-2 OS samples contain highest level of variability in $|\mu_T - TCS|$. **B.** After increasing coverage filter ±3kb around $\mu_T$ (from 0.001x, to 0.0075x), $|\mu_T - TCS|$ values between U-2 OS replicates become more reproducible and contain less variability.

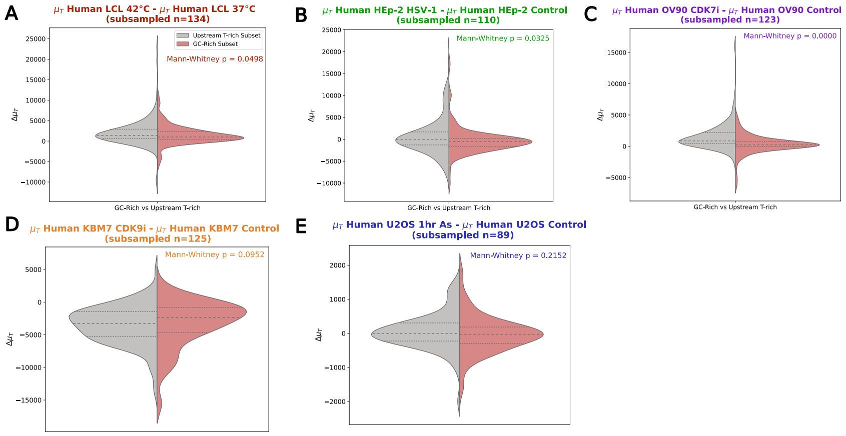


Figure S17: **Difference in run-on between upstream T-rich and GC-rich genes.** Representative distributions for subsampled gene sets (upstream T-rich subset randomly subsampled to match GC-rich subset size), sampled 4 times to control for subset size differences. Samples include **A.** heat-shock, **B.** HSV-1, **C.** CDK7 inhibition, **D.** CDK9 inhibition, and **E.** Arsenic. A Mann-Whitney U test was ran on each comparison, the highest p-values obtained are reported.

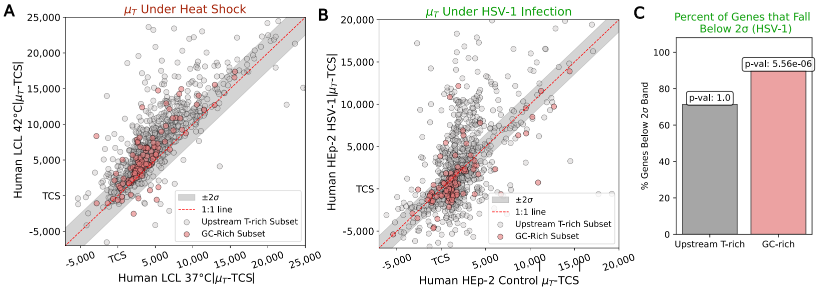


Figure S18: $|\mu_T - TCS|$ **scatter plots colored on subsets of genes (GC-rich vs. upstream T-rich genes). A.** Heat-shock and **B.** HSV-1. **C.** Hypergeometric test shows significant enrichment of GC-rich genes below $2\sigma$ bounds under HSV-1 infection.

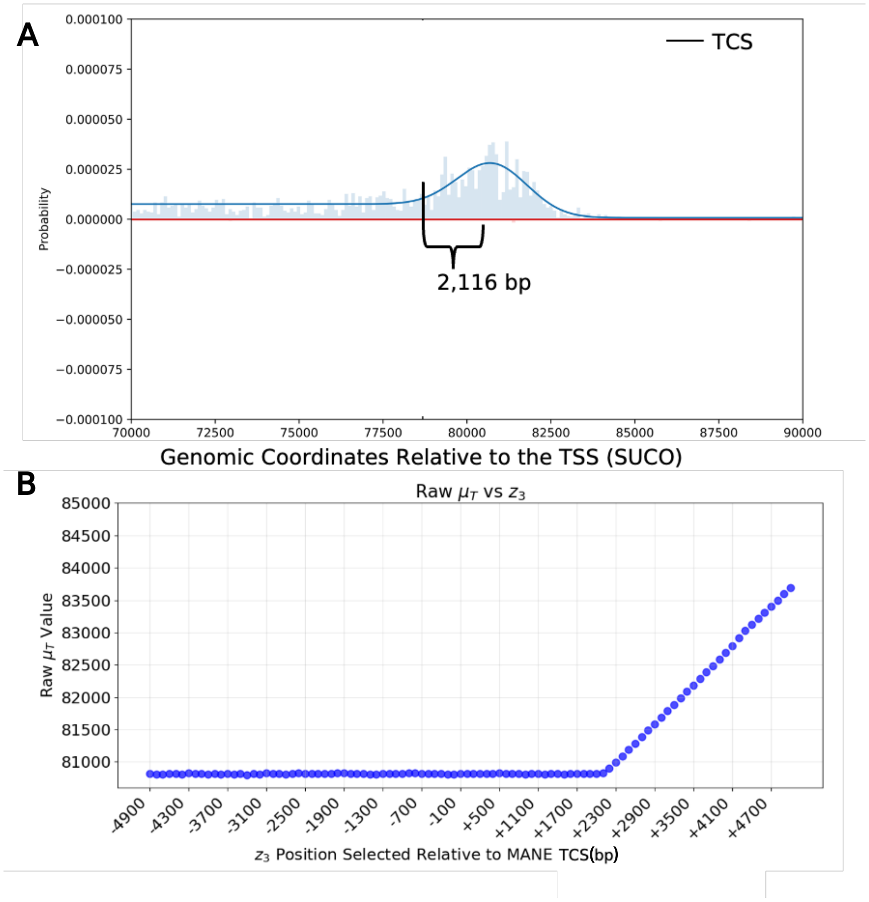


Figure S19: **Choice of $z_3$ prior does not impact where $\mu_T$ is called by LIET.** We moved the position of $z_3$ ($3'$ annotation fed into LIET[3]) in 100 base increments from -5000 to +5000 bps of the annotated MANE [4] TCS position at the gene SUCO. **A.** $3'$ end of the gene SUCO. **B.** $\mu_T$ is not impacted by the position of $z_3$ unless $z_3$ is farther downstream than the disassociation peak position.

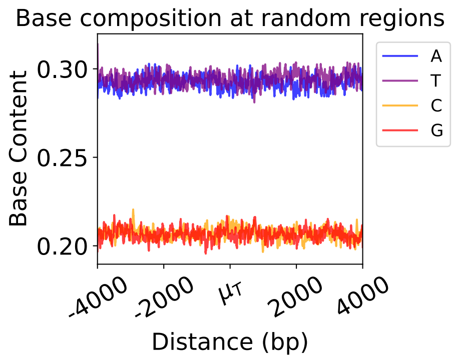


Figure S20: **Base composition at 8kb sized random regions in the human genome (hg38).** Base composition averaged across a set of random regions drawn from the human genome (hg38).

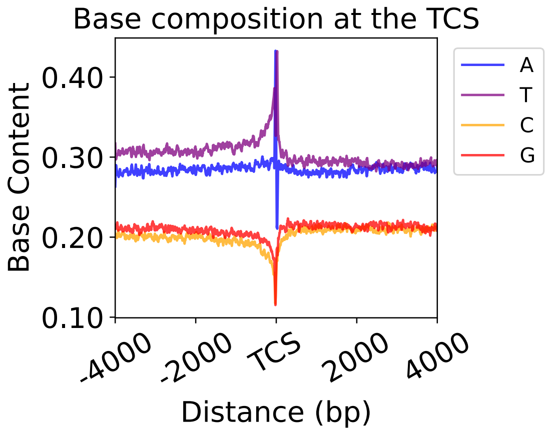


Figure S21: **Base composition ±4kb around the TCS.** T content is elevated upstream of the TCS and spikes at the TCS. A content follows a similar trend as T, but remains slightly below T content. Because of the averaging effect and variable length between the TCS and $\mu_T$, the T rich region upstream of $\mu_T$ is difficult to discern in this meta-gene.

### Supplementary Table

For the control condition analysis (Figures 1 - 3), each cell type meta-sample was generated by combining the collection of samples listed in the Accessions column of Table 1. For the perturbation analysis (Figures 4 - 5), control replicates were combined to make a meta-sample, and the same process was repeated with perturbation replicates. Meta samples in both basal and perturbed conditions were made to increase depth and complexity of the data fed into LIET, enabling LIET to output more accurate parameters. Individual control replicates were used to calculate the $2\sigma$ bounds to quantify and test changes in disassociation in Figures 4 - 5. Additionally, individual replicates were used to calculate $2\sigma$ bounds in Figure 3 D, which are $2\sigma$ bounds from the human replicates (human had a larger level of replicate variation compared to rhesus, thus was applied to Figure 3 D to be conservative).

Table 1: **Data utilized within this paper.** This analysis was performed across 9 cell types, 5 species, and 7 different perturbations.

| **Cell line** | **Tissue** | **Condition** | **Accessions** | **Species, Genome** | **Assay** |
|---|---|---|---|---|---|
| HCT116 | colorectal epithelial | control | SRR6290501<br>SRR6290500<br>SRR6290499<br>SRR9304731<br>SRR9304730<br>SRR6290518<br>SRR6290517<br>SRR6290516<br>SRR6290515<br>SRR6290510<br>SRR6290509<br>SRR6290508<br>SRR6290507<br>SRR6290502<br>SRR8867628<br>SRR6290526<br>SRR6290525<br>SRR6290523<br>SRR6290524<br>SRR8867629<br>SRR8867637<br>SRR8867636 | Human, hg38 | PRO-seq |
| HEK293/T | kidney epithelial | control | SRR6026781<br>SRR6026782<br>SRR6927839<br>SRR7988488<br>SRR7988490 | Human, hg38 | PRO-seq |

| | | | | | |
|---|---|---|---|---|---|
| K562 | bone marrow | control | SRR12083665<br>SRR8137173<br>SRR4454568<br>SRR4454567<br>SRR11793826<br>SRR11793825<br>SRR12083664<br>SRR1554311<br>SRR1554312<br>SRR5364303<br>SRR5364304<br>SRR8669163<br>SRR8669162 | Human, hg38 | PRO-seq |
| KBM7 | bone marrow | control | SRR10354625<br>SRR10354624<br>SRR10354600<br>SRR10354601<br>SRR10354602<br>SRR10354603<br>SRR10354608<br>SRR10354609<br>SRR10354610<br>SRR10354611 | Human, hg38 | PRO-seq |
| KBM7 | blood | control, CDK9i (NVP-2) | SRZ10354600<br>SRZ10354602<br>SRZ10354616<br>SRZ10354618 | Human, hg38 | PRO-seq |

| | | | | | |
|---|---|---|---|---|---|
| LCL | blood | control | SRR6727941 SRR6727942 SRR6727943 SRR6727944 SRR6727945 SRR6727946 SRR6727947 SRR6727948 SRR6727949 SRR6727950 SRR6727951 SRR6727952 SRR6727953 SRR6727954 SRR6727955 SRR6727956 SRR6727957 SRR6727958 SRR6727959 | Human, hg38 | PRO-seq |
| LCL | blood | control, heat-shock (37°C and 42°C) | SRR14352131 SRR14352132 SRR14352135 SRR14352136 | Human, hg38 | PRO-seq |
| HCT116 | colorectal epithelial | control, CDK8i (CDK8 analog sensitive, 3MB) | SRR8867628 SRR8867629 SRR8867622 SRR8867623 | Human, hg38 | PRO-seq |
| HCT116 | blood | control, IFN-$\gamma$ | SRR7824015 SRR7824016 | Human, hg38 | PRO-seq |
| U-2 OS | bone | control, arsenic exposure | GSM9321236 GSM9321237 GSM9321238 GSM9321239 | Human, hg38 | PRO-seq |

| | | | | | |
|---|---|---|---|---|---|
| OV90 | ovary | control, CDK7i (SY5609) | SRR23989399 SRR23989398 SRR23989397 SRR23989396 | Human, hg38 | PRO-seq |
| HEp-2 | larynx | mock infection, HSV-1 infection | SRR6216708 SRR6216709 SRR6216710 SRR6216711 SRR6216712 SRR6216713 | Human, hg38 | PRO-seq |
| BEAS-2B | lung/bronchus epithelial | control, wood smoke | SRR13772348 SRR13772349 SRR13772350 SRR13772351 SRR13772352 SRR13772353 | Human, hg38 | PRO-seq |
| LCL | blood | control (prepped for comparison to primate data) | GSM6716757 GSM6716758 | Human, hg38 | PRO-seq |
| LCL | blood | control | GSM6716755 GSM6716756 | Gibbon, nomLeu3 [5] | PRO-seq |
| LCL | blood | control | GSM6716759 GSM6716760 | Rhesus, rheMac10 [6] | PRO-seq |
| LCL | blood | control | GSM6716761 GSM6716762 | Squirrel, saiBol1 [5] | PRO-seq |
| ESC (strain: 129/Sv) | blastocyst | control | SRR9007616 | Mouse, mm10 [6] | PRO-seq |
| LCL | blood | control | GSM5269423 | Human, hg38 | ATAC-seq |
| HEK293 | colon | control | SRR28504065 | Human, hg38 | ChIP-seq |

### References


[1] Magda Kopczyńska, Upasana Saha, Anastasiia Romanenko, Takayuki Nojima, Michał R. Gdula, and Kinga Kamieniarz-Gdula. Defining gene ends: Rna polymerase ii ctd threonine 4 phosphorylation marks transcription termination regions genome-wide. *Nucleic Acids Research*, 53(2):gkae1240, 2025.

[2] Jean-Pierre Etchegaray, Lei Zhong, Catherine Li, Telmo Henriques, Eileen Ablondi, Tomoyoshi Nakadai, Capucine Van Rechem, et al. The histone deacetylase sirt6 restrains transcription elongation via promoter-proximal pausing. *Molecular Cell*, 75(4):683–699, 2019.

[3] Jacob T. Stanley, Georgia E. F. Barone, Hope A. Townsend, Rutendo F. Sigauke, Mary A. Allen, and Robin D. Dowell. LIET model: capturing the kinetics of RNA polymerase from loading to termination. *Nucleic Acids Res*, 53(7):gkaf246, 2025.

[4] Julio Morales, Shweta Pujar, Jane E. Loveland, Andrey Astashyn, Rosie Bennett, Ann Berry, Eleanor Cox, Chris Davidson, Olga Ermolaeva, Catherine M. Farrell, and Rabiya Fatima. A joint ncbi and embl-ebi transcript set for clinical genomics and research. *Nature*, 604(7905):310–315, 2022.

[5] Zoonomia Consortium. A comparative genomics multitool for scientific discovery and conservation. *Nature*, 587:240–245, 2020.

[6] L. F. K. Kuderna, J. C. Ulirsch, S. Rashid, et al. Identification of constrained sequence elements across 239 primate genomes. *Nature*, 625:735–742, 2024.
